# Printability study of metal ion crosslinked PEG-catechol based inks

**DOI:** 10.1101/599290

**Authors:** Małgorzata K. Włodarczyk-Biegun, Julieta I. Paez, Maria Villiou, Jun Feng, Aranzazu del Campo

## Abstract

Inspired by reversible networks present in nature, we have explored the printability of catechol functionalized polyethylene glycol (PEG) based inks with metal-coordination crosslinking. Material formulations containing Al^3+^, Fe^3+^ or V^3+^ as crosslinking ions were tested. The printability and shape fidelity were dependent on the ink composition (metal ion type, pH, PEG molecular weight) and printing parameters (extrusion pressure and printing speed). The relaxation time, recovery rate and viscosity of the inks were analyzed in rheology studies and correlated with thermodynamic and ligand exchange kinetic constants of the dynamic bonds and the printing performance (i.e. shape fidelity of the printed structures). The relevance of the relaxation time and ligand exchange kinetics for printability was demonstrated. Cells seeded on the crosslinked materials were viable, indicating the potential of the formulations to be used as inks for cell encapsulation. The proposed dynamic ink design offers significant flexibility for 3D (bio)printing, and enables straightforward adjustment of the printable formulation to meet application-specific needs.

## 1. Introduction

Hydrogels are commonly used in 3D extrusion bioprinting. [1] They allow to mimic the mechanical properties and hydration of biological tissues and show reasonable biocompatibility for cell encapsulation. [2] However, a typical hydrogel formulation used for cell encapsulation is not necessarily printable. [2, 3] A good hydrogel for 3D bioprinting needs to be easily extruded and quickly solidify upon deposition. These abilities determine the accuracy and reproducibility of the printed structures, the time required for printing and the resolution of the printed object. [1] Additionally, the required printing parameters (velocity, pressure), crosslinking chemistry and precursors should be compatible with living cells. Reactive hydrogels used for 3D printing make use of crosslinking agents [2, 4] or photo-polymerization of acrylate functional groups [4] for fast gelation. However, mixtures of fast reactive species typically result in narrow printability windows and low processability [3] due to limited control over the kinetics of the crosslinking reaction [5]. Fast reactions lead to inhomogeneity of the printed structures or cell damage due to the need of high shear forces during printing. Slow kinetics lead to prints with poor shape fidelity and sedimentation of cells in the printed structures. [4] On the other hand, photopolymerization after or during printing might be harmful to some cell lines [6] and leads to uneven irradiation doses in consecutively printed layers.

A promising approach for printing hydrogels is the use of networks with reversible bonds. Reported examples make use of ionic interactions (eg. alginate [7]), dynamic covalent bonds (eg. hydrazone crosslinks [8]), guest-host complexes [9], hydrogen bonds [10] or metal-ligand coordination complexes [3, 11]. Networks with reversible bonds have several advantages over non-reversible systems for printing: they possess inherent shear thinning and self-healing properties; allow processability independent of gelation kinetics; do not require exposure to harsh conditions (eg. UV irradiation or reactive crosslinking agents); do not need addition of thickening agents to increase viscosity to facilitate extrusion. [4] Among reversible bonds, metal-ligand coordination complexes offer wide versatility in terms of bonding strength and bonding dynamics by varying the metal ion or the pH of the system [3], leading to networks with adjustable mechanical properties [11]. Metal complexation is widely used in natural materials to tune mechanical strength of bulk and interface materials. [3] One of the most prominent examples is the metal-catechol interaction in the DOPA-reach mussel adhesive proteins. The catechol-metal interaction takes place in seawater due to the mild basic pH and the presence of metal ions, and leads to fast gelation of the secreted proteinaceous fluid. In a similar way, catechol-functionalized polymers mixed with metal ions (V^3+^, Fe^3+^, Al^3+^) have been demonstrated to show pH-tunable crosslinking degree, fast network formation and self-healing behavior. [12, 13] The mechanical properties (e.g. shear modulus, relaxation time) of the formed polymer network can be finely tuned by the type of metal ion and pH. [13, 14]

Only a few reports have exploited metal-ligand coordination complexes to obtain printable hydrogels [3, 11, 15]. The bisphosphonate-Ca^2+^ interaction has been used to print bisphosphonate-functionalized hyaluronic acid (HA-BP) at pH 7 [11]. The reversible gel composed of 2.7% w/v HA-BP and 200 mM CaCl_2_ was printable and scaffolds with four layers, with strand diameter of ca. 1 mm, were obtained without the need of further crosslinking. The gels were stable in PBS solution at pH 7 for 1 day but softened and dissolved within hours at lower pH. This is due to the protonation of bisphosphonate at decreasing pH, and the weakening of bisphosphonate-Ca^2+^ coordination. Crosslinking with Ag^2+^ was tested as an alternative [16], however the printability was not assessed. In a different work, the carboxyl-Fe^3+^ interaction was used to stabilize poly(acrylamide-co-acrylic acid) printed structures by immersing the prints in the metal ion salt, leading to robust gel formation [3]. Strands with diameter between 0.5 – 2 mm were obtained by extrusion of a 13 % wt polymer solution. The remarkable mechanical properties of the gel originated from the high polymer content, the broad range of coordination complexes and the stabilizing ionic interactions between carboxyl and amino groups. The strong carboxyl-Fe^3+^ associations served as permanent crosslinks and provided gel integrity and high stiffness, whereas the weaker complexes served as reversible bonds, dissipating energy, and contributing to gel toughness. The influence of acrylic acid unit density, metal ion type and concentration on gel formation was tested. Crosslinking with Al^3+^, Mg^2+^, Ca^2+^, Zn^2+^, Ni^2+^ was attempted, but only mixing with Al^3+^ and Fe^3+^ led to gel formation, with significantly stronger gels obtained by crosslinking with Fe^3+^ ions. This effect was assigned to the highest coordination strength and stability of the carboxyl-Fe^3+^ interaction. Finally, the influence of pH on the coordination state and stability constant of the complexes was investigated as a tool to introduce pH-driven tunability of mechanical properties, shape memory properties and sol-gel transition. Incorporation of sodium alginate (SA) to the polymer mixture led to printed polymers with improved mechanical stability [15] where Fe^3+^ ions crosslinked with the carboxyl groups from both polymers. However, in order to achieve printable formulations, 10% silicone had to be added to the SA/P(AAm-co-AAc) mixture in order to obtain viscous prepolymer mixtures that could be extruded. In addition, a material deposition had to be followed by 10 minutes UV irradiation, 3 h of soaking in Fe(NO_3_)_3_·9H_2_O solution, and 48 h incubation in water to remove excess of Fe^3+^ ions. With the proposed system prints with resolution limited to a strand size of ~800 μm were obtained. However, the multiple and long processing steps make the system unpracticable from an application point of view.

Here, we present printable polyethylene glycol (PEG) based formulations with catechol-metal complexes as reversible crosslinks able to support network formation at close-to-physiological pH and in the absence of harsh agents. The catechol-metal bonds serve as temporary crosslinks during printing, with possibility of post-printing stabilization by gentle oxidation to form a covalent network. The inks can be printed without the use of a supportive bath or any additives to tune viscosity. No UV post-printing treatment is necessary to further stabilize the printed structures. In addition, the reactivity of catechol groups allows effective fixing of the printed material to different surfaces, including natural surfaces or tissues, eventually in wet conditions [17]. This system complements the recent report by Burdick et al. [10] utilizing gallols for short-term gelation via reversible hydrogen bonding, and slow, spontaneous oxidation for long-term stabilization of the printed scaffolds. Distinctively, the metal complexation approach in our system offers flexible and controlled tuning of viscosity and mechanical properties, and adjustable ink formulation for specific application demands, including possibility of using different catechol-functionalized polymeric materials. Additionally, printing can be performed with broad range of printing parameters (speeds and pressures) and smooth strands with distinctively small diameters can be obtained.

This article reports on the rheological properties and printability window of PEG-catechol based inks crosslinked with V^3+^, Fe^3+^, Al^3+^ ions, highlighting possible relationships between rheological parameters, thermodynamic and kinetic properties of the metal-catechol bond and printing results at different printing conditions. The study complements current efforts to correlate rheological behavior and printability of hydrogel-based materials [18–23], and contributes to the progress in the design and formulation of advanced inks.

## 2. Materials and Methods

### 2.1 Materials

4-arm PEG succinimidyl carboxymethyl ester (PEG-NHS, MW of 10, and 20 kDa) was purchased from JenKem Technology (TX, USA). All other chemicals were purchased from Sigma-Aldrich (Germany), unless stated differently. Printing needles were provided by Vieweg (German), Optimum^®^ cartridges and pistons for printing by Nordson (Germany).

### 2.2 Synthesis of PEG-Dopamine

Dopamine-functionalized PEG (PEG-Dop) was synthesized according to a previously reported protocol [5, 17, 24]. 110 μL of N-methylmorpholine (1.0 mmol) and 114 mg of dopamine (0.6 mmol) were mixed in 5.0 mL of anhydrous dimethylformamide (DMF) with magnetic stirring in argon atmosphere for 20 min. 1g (0.1 mmol) of 4-arm PEG succinimidyl carboxymethyl ester (MW: 10 kDa or 20 kDa) was dissolved in 5.0 mL of anhydrous DMF and was added dropwise into above mixture. The reaction was performed for 24 hours at room temperature under Ar atmosphere, with constant stirring. After evaporation of DMF at reduced pressure, the crude product was redissolved in deionized water (pH 6.0) and dialized against deionized water (pH 6.0) with membrane tubing (MW cut-off of 3.5 kDa) for 2 days (dialysate was changed three times per day). The purified product was freeze dried and stored under −20 °C until use. The obtained PEG-Dop was characterized by 1H-NMR (Bruker Advance III HD, 300MHz/54mm, Fällanden, Switzerland) in D2O to check the extent of the reaction and calculate the degree of catechol substitution. This was calculated by comparing the integral of the signal of PEG backbone (3.70-3.50 ppm which integrates for 210H or 437H for PEG with molecular weight 10 or 20 kDa, respectively) to the integral of the aromatic protons of dopamine (6.80-6.50 ppm, 3H). The obtained catechol substitution degree in all PEG batches was >85%.

### 2.3 Ink formulation

A 10% (w/v) solution of PEG-Dop (10 kDa or 20kDa) in Milli-Q water was prepared and vortexed to ensure homogeneous mixing. Metal salt solutions (FeCl_3_, AlCl_3_ and VCl_3_) were prepared in Milli-Q water at 40 mM concentration. The inks were prepared by mixing 1/2 final volume of PEG-Dop solution with 1/6 final volume of metal ion solution and, in the last step, 1/3 final volume of NaOH base (molarity indicated in Table 1) in order to achieve homogenous cohesive networks. For example, to prepare 750 μl of V^3+^ crosslinked PEG-Dop ink: 375 μl of 10% (w/v) PEG-Dop, 125 μl of 40 mM VCl_3_ and 250 μl of 100 mM NaOH were used. The sample was stirred with spatula in the Eppendorf tube until a homogenous sample was obtained. The final concentration of PEG-Dop in the samples was 5%, and the molar ratio of dopamine to metal ions was 3:1. The pH of crosslinked network was measured with a pH-meter with flat surface electrode (PH100 Waterproof ExStik^®^, Extech Instruments, USA). All the ink formulations used in this study are presented in Table 1.

**Table 1.** Composition and final pH of all the ink formulations used in the study.

| Ink abbreviation | MW of PEG-Dop | Crosslinker | Final NaOH concentration | Final pH |
| --- | --- | --- | --- | --- |
| **PEG-Dop/Al<sup>3+</sup>** | **10 kDa** | **AlCl<sub>3</sub>** | **33 mM** | **8.9** |
| PEG-Dop/Al<sup>3+</sup> (pH 8.4) | 10 kDa | AlCl<sub>3</sub> | 30 mM | 8.4 |
| **PEG-Dop/Fe<sup>3+</sup>** | **10 kDa** | **FeCl<sub>3</sub>** | **33 mM** | **8.9** |
| PEG-Dop/Fe<sup>3+</sup> (20 kDa) | 20 kDa | FeCl<sub>3</sub> | 15 mM | 8.6 |
| **PEG-Dop/V<sup>3+</sup>** | **10 kDa** | **VCl<sub>3</sub>** | **33 mM** | **8.6** |

\* Main samples discussed in the study indicated in bold. For detailed results obtained for PEG-Dop/Al<sup>3+</sup> (pH 8.4) and PEG-Dop/Fe<sup>3+</sup> (20 kDa) inks see Supplementary Information.

### 2.4 Rheological characterization

The rheological properties of PEG-Dop networks crosslinked with different metal ions (PEG-Dop/M^3+^) were measured using a rotational rheometer (DHR3, TA Instruments, USA) equipped with a 8 mm parallel plate geometry. A material volume of 8 μl was used for the measurements. A measuring gap of 200 μm was set and the experiments were performed at controlled temperature of 23°C. To avoid drying of the sample during testing, paraffin oil was applied around the samples after setting measuring gap and evaporation blocking cap was installed. All rheological experiments were performed at least in triplicate. For frequency test, 2 of 6 measurements conducted for PEG-Dop/Al^3+^ were excluded from the final data set, due to the one order of magnitude longer relaxation times than achieved in remaining 4 measurements.

#### 2.4.1 Characterization of time dependent response and final stiffness

Frequency sweep oscillatory test was performed at 1% strain. The storage (G’) and loss modulus (G”) were recorded over the frequency range of 10^−2^ to 10^2^ Hz. The crossover modulus was taken as the modulus value at which G’ intersects G”. The relaxation time τ corresponded to the reciprocal of the frequency at which G’ intersects G”. The final material stiffness was analyzed by measuring G’ and G” as a function of time in an oscillatory test at a constant frequency (f = 10 Hz) and strain (γ = 1 %). After 180 s, the gels were broken (f = 10 Hz and γ = 0.01 to 1000 %) to monitor self-healing behavior. Subsequent gel re-formation was analyzed for 1 h at constant frequency (f = 10 Hz) and strain (γ = 1 %). All tests were initiated 20 minutes after sample loading to the rheometer.

#### 2.4.2 Characterization of shear thinning

The viscosity measurements were performed in the rotational flow sweep experiments. The same samples were used as for the frequency sweep oscillatory test. After 20 minutes of a soak time (to heal after oscillatory test), viscosity was recorder during the increasing shear rate *ẏ* = 0,001 · 1/s to 3000 · 1/s (shear rate given at the edge of the rotating plate).

### 2.5 Evaluation of printability

#### 2.5.1 Filaments printing and characterization

PEG-metal inks were printed using pneumatic extrusion-based BioScaffolder 3.1 (GeSiM, Germany). For all experiments a 25 G conical nozzle (inner diameter: 260 μm) was used. Extrusion pressure was in the range of 15 - 550 kPa (18 different levels, Table S4) and the speed of printing head was set to 1, 2, 5 or 10 mm/s. The inks were prepared freshly before printing, loaded to 10 ml cartridges, closed with pistons to prevent leaking and ensure laminar ink flow, and after 20 minutes waiting time printed at room temperature into polystyrene 6-well cell culture plate placed on the holder on the printing stage. Each structure was imaged immediately after printing with stereomicroscope SMZ800N (Nikon, Germany) with home-made bottom illumination. Strand width was measured (9 measurements per sample) with the integrated NIS-Elements D (Nikon, German) software.

#### 2.5.2 Printing in 3D

Crosslinking solution was prepared by mixing 50 ml of 36 mM sodium periodate (NaIO_4_) with 30 ml MiliQ water and 20 ml 100 mM NaOH, to obtain an oxidizing solution with final oxidant concentrations of 18 mM NaIO_4_ and pH ~ 9.

PEG-Dop 3-dimensional multilayer grid-like constructs were printed based on the square design with 1 cm x 1 cm dimensions (for PEG-Dop/V^3+^) or 9.1 cm x 9.1 cm (PEG-Dop/Al^3+^ and PEG-Dop/Fe^3+^), and consecutive layers rotated by 90°. After printing of each layer, ca. 2 ml of crosslinking solution was applied for 30 seconds on top of the print to introduce covalent crosslinking, and removed before printing of the next layer. For a PEG-Dop/Al^3+^, 1-4 layers constructs were printed with 85 kPa pressure and 3 mm/s printing speed. For a PEG-Dop/Fe^3+^, 1^st^ layer was printed with 50 kPa pressure and 2 mm/s printing speed, all the consecutive layers were printed with 100 kPa pressure and 2 mm/s printing speed. For PEG-Dop/V^3+^, 1-layer construct was printed with 200 kPa pressure and 2 mm/s printing speed, 2-4 layers constructs were printed with 250 kPa pressure and 1 mm/s printing speed.

### 2.6 Cell viability studies

#### 2.6.1 Cell culture

L929 Fibroblasts (ATCC, Germany) were cultured in RPMI 1640 medium (Gibco, 61870-010) supplemented with 10 % FBS (Gibco, 10270), 200 mM L-glutamine and 1% penicillin/streptomycin (Invitrogen), at 37 °C in a humidified atmosphere of 5 % CO_2_. Cell culture media was changed every second day. Cells from passage 40-45 were used.

#### 2.6.2 Live/dead assay

PEG-Dop/M^3+^ mixtures were prepared as described in section 2.3. Samples of 30 ul of each ink type were molded into the 96-well. Half of the samples was used without subsequent treatment (metal-crosslinked samples), half was covered with 150 ul crosslinking solution (composition as in the section 2.5.2) for ca. 15 minutes (covalently crosslinked samples). All samples were washed with ~ 200 ul of medium prior to cell seeding. Samples were prepared in triplicates, with exception of PEG-Dop/Al^3+^ at pH 8.9 (duplicate).

200 ul of medium containing 25 000 cells was deposited on top of each sample. The samples were incubated for 1 h at 37 °C in a humidified atmosphere of 5% CO_2_ and cell viability was assessed. Staining protocol with a fluorescein diacetate (FDA) to detect live cells and propidium iodide (PI) to detect dead cells, was used as follows. 1 μg/ml PI solution and 1 μg/ml FDA solution were dissolved in PBS to achieve final concentrations of 20 μg/ml and 6 μg/ml, respectively. Cell culture medium was removed from the samples, and 100 μl of staining solution was added to each well for 10 min. Samples were 2 x washed with PBS and imaged with PolScope microscope (Zeiss, Germany). Image analysis was performed with ImageJ software. After subtracting background (rolling ball radius: 100) counting of cells was performed with the function “find maxima” (noise tolerance: 100 for red channel and 1000 for green channel).

### 2.7 Statistical analysis

All the results are reported as the mean ± standard deviation. Statistical differences were analyzed based on one-way Analysis of Variance (ANOVA) performed using Excel Data Analysis or In Stat3 software. Differences for p < 0.05 were considered as significant.

## 3. Results

### 3.1 Ink development

PEG-Dop was mixed with the solutions of metal ions: Al^3+^, Fe^3+^, V^3+^. These cations form metal-catechol complexes with comparable thermodynamic binding constant [25, 26] but different water exchange rate [27] and coordination degree at pH ~8.0 [14] (see Table 2 for comparative values). These differences are expected to affect the flow properties of the solutions [28]. The molar ratio of dopamine groups to metal ions was 3:1 in order to maximize the coordination degree of the complex. Since the solubility of metal cations is low at basic pH [13], the solutions of PEG-Dop polymer and cations were first mixed at acidic pH (pH 3-4) and then the pH was increased to pH ~8.5 to allow complex formation. The immediate increase in viscosity at basic pH (and color change to purple and dark blue for PEG-Dop/Fe^3+^ and PEG-Dop/V^3+^, respectively) indicated the fast formation of coordination bonds between the ionized catechol end-groups of the PEG-Dop chains and the metal ions [25, 29]. PEG-Dop/V^3+^ formed crosslinked network almost instantly, while PEG-Dop/Fe^3+^ and PEG-Dop/Al^3+^ required longer time (in 10 – 20 s range) to form homogenous, viscous mixtures (slightly longer time for PEG-Dop/Al^3+^ than PEG-Dop/Fe^3+^). Preliminary screening of PEG-Dop/Fe^3+^ mixtures at different pH (pH 2.5 – 12.5) and polymer concentrations (2.5 %, 5 % and 10 % (w/v)) was performed in order to identify suitable ink compositions for extrusion printing. 5 % PEG-Dop (10kDa) solutions at pH 8.6 – 8.9 gave the best results in terms of ink homogeneity and handling (for more details see Supplementary Information Table S1 and Fig. S1). Table 1 presents the selected compositions from the preliminary screening, used for the printing experiments in the following sections.

**Table 2.** Thermodynamic and kinetic parameters of catechol-metal coordination complexes: degree of coordination at pH 8.0, metal-catechol binding constant in tris-coordinated complexes, and water exchange rate constant.

| Metal | Degree of coordination [14] | Log $\beta_3$ | Water exchange rate constant: $k$ [27] |
| --- | --- | --- | --- |
| $Al^{3+}$ | Mono | 41 * | 1.92 |
| $Fe^{3+}$ | Bis | 46,5* / 41** | $1.6 \times 10^2$ |
| $V^{3+}$ | Tris | 40,5** | $5 \times 10^2$ |

\* Binding constant from [25], at 0.1 M KCl ionic strength and 25 °C
\*\* Cumulative binding constant calculated based on [26]

### 3.2 Rheological evaluation of the PEG-Dop/M^3+^ inks

The rheological properties of the material determine its suitability for printing. Frequency sweep measurements revealed liquid-like behavior of PEG-Dop inks crosslinked with Al^3+^ and Fe^3+^, and predominantly solid-like behavior of PEG-Dop/V^3^ networks between 10^−2^ and 25 Hz (Fig. 2A). The G’ and G’’ curves crossed at 4.3, 5.5 and 0.025 Hz in PEG-Dop inks crosslinked with Al^3+^, Fe^3+^ or V^3+^ respectively. These frequencies define the relaxation time of the networks, i.e. the time required to release an applied stress. Fe^3+^ and Al^3+^ crosslinked networks showed comparable relaxation times at pH 8.9, in the order of 0.2 second (Table 3). The relaxation time did not change with the molecular weight of the PEG-Dop (Table S2), and was shorter at lower pH (τ = 0,12 s for PEG-Dop/Al^3+^ (pH 8.4), Table S2). The PEG-Dop-V^3+^ network showed longest relaxation time (τ = 44 s at pH 8.6). Previous studies have found relaxation times of ~0.09 s, 0.41 s and 7.6 s for 10 kDa 4-arm PEG-Dop hydrogel, with ~15% (w/w) polymer concentration, crosslinked with Al^3+^, Fe^3+^ and V^3+^, respectively, at pH 8.0. [14] The relaxation time is expected to depend on: (i) the kinetics of ligand exchange of the metal-ligand coordination bond [30], and (ii) the stoichiometry of the coordination complex, which relates to the ionization state of the catechol and therefore to the pH of the solution [14]. UV-Vis spectroscopy studies showed that PEG-Dop at pH 8.0 forms predominantly tris-coordinated complexes with V^3+^, bis-coordinated complexes with Fe^3+^, and mono-coordination complexes with Al^3+^. [14] Note that in the context of printing application, longer relaxation times imply that network rearrangements under applied strain occur at slower rate [16], and this hampers the extrusion process. On the other hand, long relaxation times lead to better shape fidelity after 3D deposition [31].

**Figure.1.**
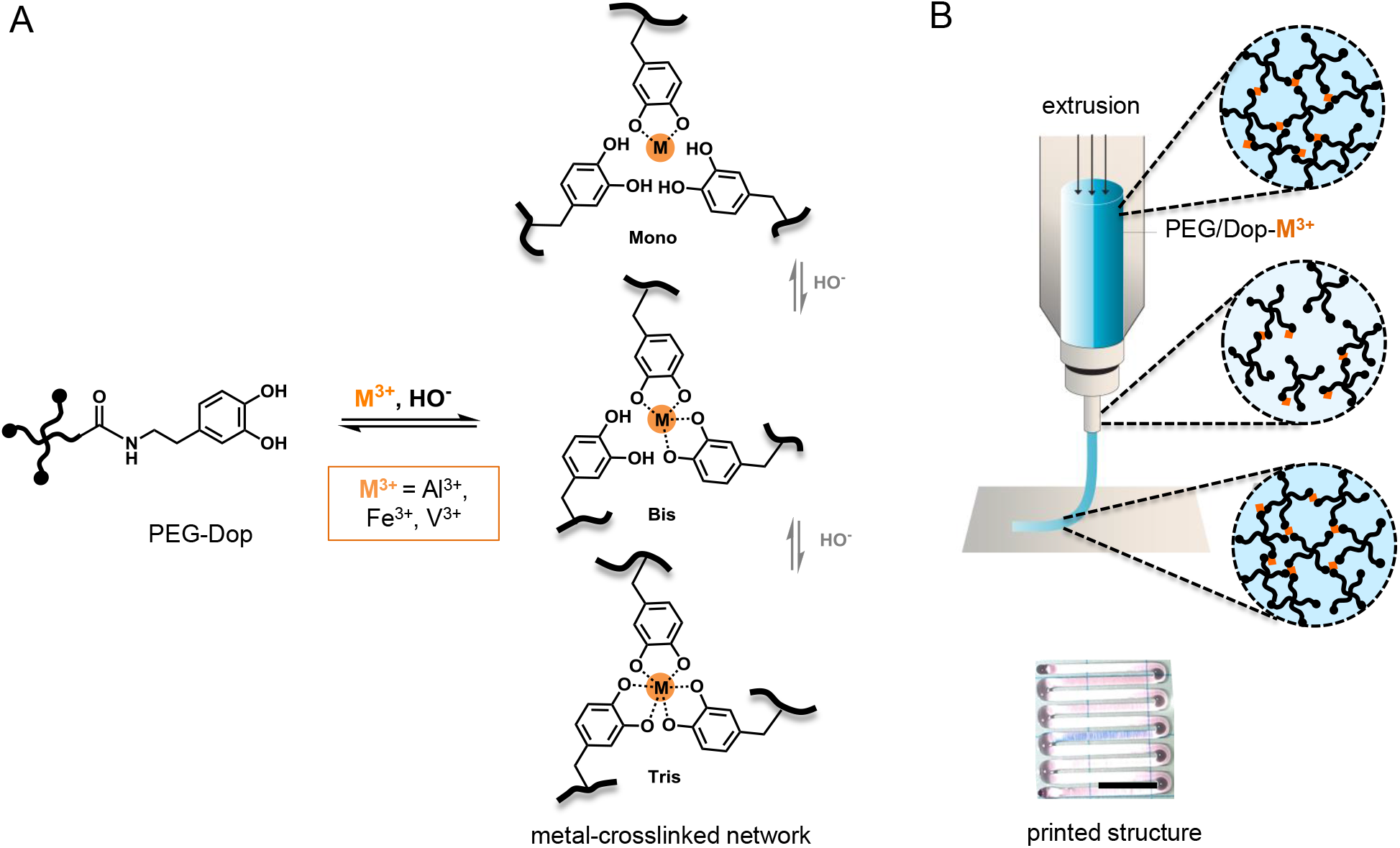
**A.** Schematic of metal-ligand complexes formed by PEG-Dop/M^3+^. The coordination number (mono-, bis-, tris-) varies with the coordinating metal type and the ionization state of the catechol, influenced by pH. **B.** Extrusion process of PEG-Dop/M^3+^ ink with indicated network rearrangement under applied shear force due to the dynamic character of the bonds, and camera image of a printed PEG-Dop/Fe^3+^ strand. Scale bar: 500 μm.

**Figure.2.**
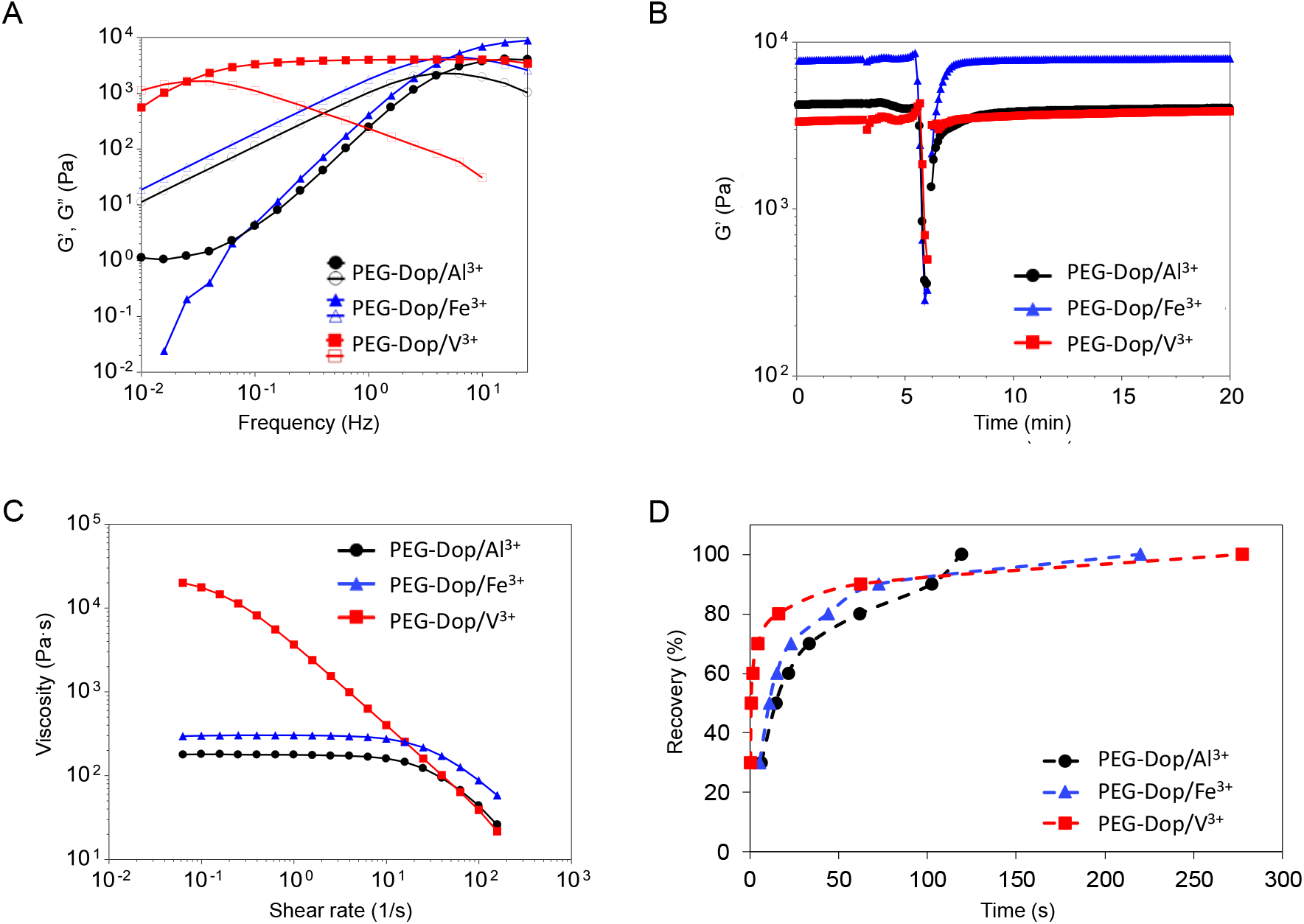
Rheological curves of PEG-Dop networks with metal complexation crosslinking. **A.** The storage (full symbols) and loss (open symbols) modulus, G’ and G”, recorded as a function of frequency at 1% strain. **B.** Time sweep at 10 Hz frequency and 1% strain, followed by logarithmic strain increase from 0.01 to 1000% (initiated 3 after min of measurement), leading to network breakage at ~5.5 min. After 6 minutes network recovery was recorded in the consequent time sweep measurement. **C.** Viscosity of the inks as a function of shear rate, obtained from Cox-Merz rule applied to oscillatory data from graph A. **D.** Average time necessary for the recovery of the tested inks after breakage by the applied strain increasing from 0.01 to 1000%. The lines serve as a guide for the eye.

**Table 3.** Relaxation time and storage modulus of tested inks.

| Ink | Relaxation time<br>($\tau$) (s) | Storage modulus<br>($G'$) (Pa) * |
| --- | --- | --- |
| PEG-Dop/$Al^{3+}$ | $0.23 \pm 0.02$ | $4020 \pm 973$ |
| PEG-Dop/$Fe^{3+}$ | $0.20 \pm 0.06$ | $7233 \pm 1406$ |
| PEG-Dop/$V^{3+}$ | $44 \pm 19$ | $2732 \pm 1162$ |

\* Measured at 10 Hz frequency and 1% strain

The stiffness of the networks and the ability to recover after damage at high strains was characterized in time by sweep experiments at 10 Hz frequency and 1% strain (Fig. 2B). The storage modulus G’ of the network was higher for Fe^3+^ crosslinked networks (7 kPa) than for Al^3+^ and V^3+^ (4 and 3 kPa, respectively; see Table 3). This significant difference has been attributed to Fe^3+^-triggered oxidation of the catechol system and formation of covalent crosslinks in the network [14]. After breaking the network by high strains, the storage modulus dropped significantly (Fig. 2B, drop in G’ at t = 5.5 min), but the initial value of G’ was recovered within a few minutes. The healing rate was evaluated as the time required to recover a certain percentage of initial G’ value. Values are represented in Fig. 2D. PEG-Dop/V^3+^ showed the fastest healing rate, with 50% of initial G’ value recovered in less than one second. PEG-Dop/Al^3+^ and PEG-Dop/Fe^3+^ showed 10-fold longer recovery times. The observed trend in the recovery time PEG-Dop/V^3+^>> PEG-Dop/Al^3+^≈ PEG-Dop/Fe ^3+^ is expected to be related to the kinetic constant of the metal-ligand bond [29]. The water exchange kinetic constant decreases in the order V^3+^ > Fe^3+^ >> Al^3+^ (Table 2), suggesting that this parameter explains only in part the observed recovery behavior. Note that the ability of the PEG-Dop/M^3+^ system to quickly reform the crosslinking points makes it an interesting candidate for printable materials. During the extrusion process material has to flow through the nozzle and needs to undergo rapid re(gelation) just after ejection in order to maintain high shape fidelity of the printed structure [4].

The shear thinning behavior of the inks was investigated during continuous rotation by monitoring the viscosity changes at increasing shear rate (Fig. S3A). However, measurements in rotational mode resulted in edge failure and material escape from the rheometer plates (see Supplementary Information, Fig. S3B). Therefore, the Cox-Merz rule was applied to the oscillatory data at small oscillations, as no escape of the material occurred (Fig. 2C). The Cox-Merz rule is an empirical relationship which allows to predict the shear-dependent viscosity based on the oscillatory data. The shear thinning data obtained with flow experiments corresponded to the ones obtained from the Cox-Merz rule, indicating applicability of the model (see Supplementary Information, Fig. S3A). Shear thinning in Fe^3+^ and Al^3+^ crosslinked networks was observed at shear rate above 10 · 1/s (Fig. 2C). The shear thinning behavior was not affected by the molecular weight of the PEG-Dop, as tested in Fe^3+^ crosslinked polymer, or by a pH change to 8.4 in the Al^3+^ crosslinked polymer (Fig. S4A, B). The V^3+^ crosslinked polymer showed shear thinning over the whole range of shear rates investigated (10^−2^ – 10^3^ · 1/s), revealing earlier end of Newtonian plateau regime. As shown on Fig. 2C, PEG-Dop/V^3+^ was characterized by the highest zero shear viscosity (plateau observed at low shear rate), which indicates the highest resistance to deformation. The zero shear viscosity is the most commonly used parameter to characterize the bulk viscosity of shear-thinning polymeric materials. [32] For PEG-Dop/V^3+^ the zero shear viscosity was expected to appear at rates < 10 ^−1^ · 1/s (Fig. 2C and Fig. S3A), at the level of 20 kPa·s. PEG-Dop/Fe^3+^ and PEG-Dop/Al^3+^ showed 2 orders of magnitude lower zero shear viscosity values (~200 Pa·s).

Based on reported procedures [18, 33] to obtain the theoretical shear rate and the apparent viscosity while printing, we calculated maximum shear rates that PEG/Dop-M^3+^ experience during extrusion (for more details see Supplementary Information). Note that this calculation assumes that an ideal strand is printed (i.e. width of strand equal to needle diameter). Based on the calculations, maximum shear rates experienced during printing process are above 10 · 1/s. Therefore, the inks are expected to show shear thinning behavior at most of the printing conditions applied in this study (Table S3). Additionally, the inks recovered their zero shear viscosity values when measured at 1/s shear rate, imitating material at rest, after applying higher shear rates (10^2^ · 1/s, imitating material while printing)(see Fig. S4C-E). Shear thinning [2, 10] and the ability to quickly recover initial viscosity after applied high shear [18, 22], are indicators of well-printable materials.

### 3.3 Filament printing

The printability of PEG-Dop networks crosslinked with different metals ions was evaluated. We used an extrusion printer with a 260 μm nozzle, and variable printing speed (1 - 10 mm/s) and printing pressure (50 - 500 kPa). We searched for the “printable window” for each ink composition by identifying the printing parameters at which a continuous strand could be printed (including printed strands with rough or inconsistent edges). In order to characterize the strands we measured the filament width. All printed structures and the mean strand widths are presented in in Supplementary Information (Table S4.).

5% PEG-Dop formulations with Al^3+^, Fe^3+^ and V^3+^ metal ions were printable, though the printing parameters were different for each PEG-Dop/M^3+^ ink. PEG-Dop/V^3+^ was only printable within a narrow range of printing conditions (Fig. 3A, B). The printable window of PEG-Dop/Al^3+^ was significantly broader, and PEG-Dop/Fe^3+^ showed the broadest printability. Relatively high pressure was required for printing of PEG-Dop/V^3+^, presumably due to its high viscosity. The results of theoretical calculations of inks apparent viscosity and shear rates while printing (for more information see Supplementary Information and Table S3), corroborate the need of higher shear rates to extrude V^3+^ crosslinked ink.

**Figure 3.**
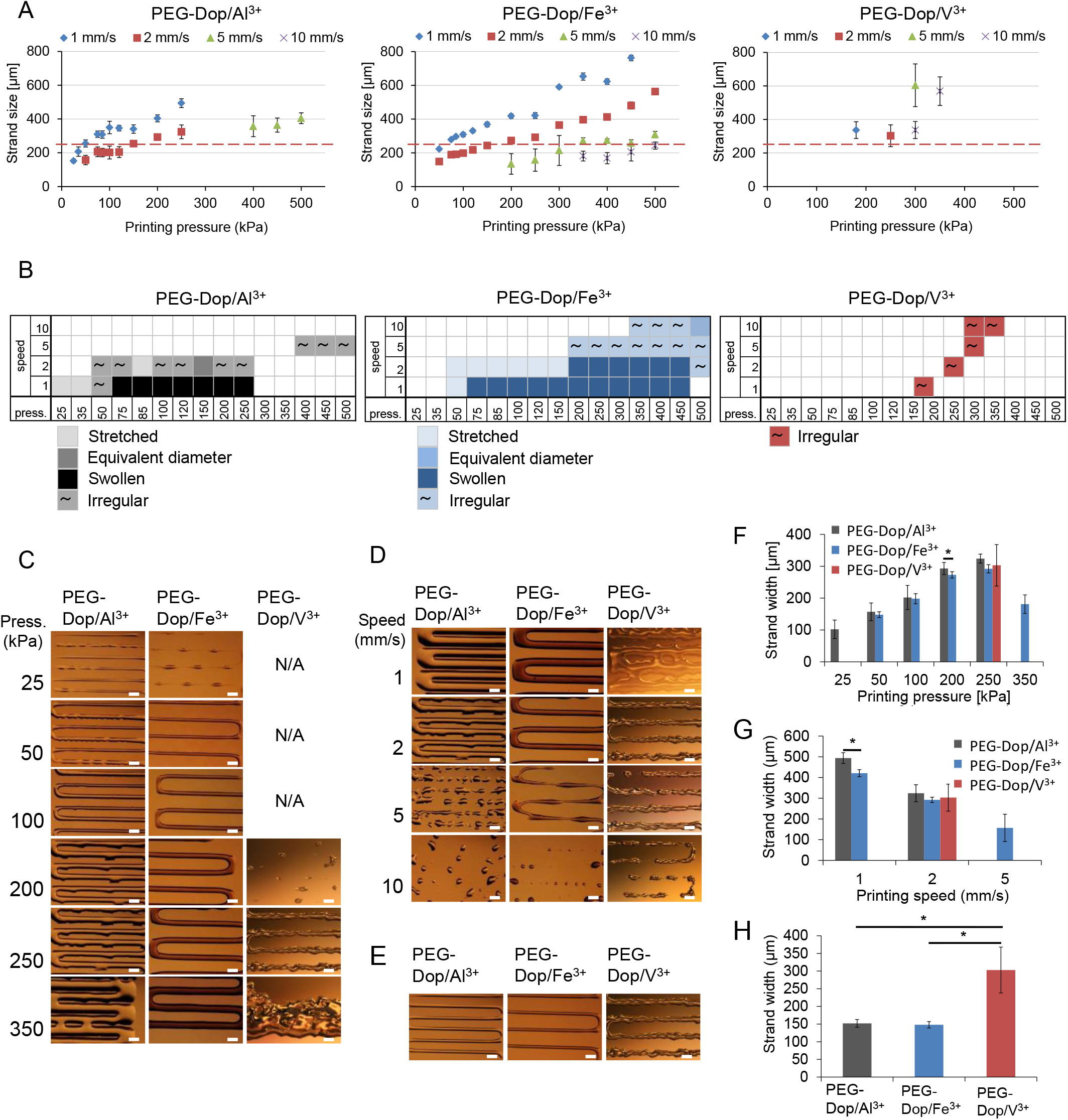
Printability of the tested inks. **A.** Strands diameter depending on head moving speed and printing pressure. Red lines serve as a guide for the eye showing the strand width value equal to the printing nozzle diameter (260 μm). **B.** Printable conditions in terms of printing pressure and speed, with marked by color shades the shape of the obtained strands according to classification by Jin et al. [34]s: irregular (~), well-defined with diameter equivalent to the diameter of the printing nozzle, swollen and stretched. **C.** Influence of pressure for printing of different inks at the head moving speed of 2 mm/s. **D.** Influence of head moving speed on printing of different inks at pressure of 250 kPa. **E.** Printed lines with the highest accuracy, defined as the thinnest printed strands with possibly smooth edges. **F. G. H.** Quantification of strand diameters corresponding to C, D, E, respectively. Scale bars: 500 μm.

The filaments printed at the different conditions were imaged and classified as: well-defined or irregular, according to the classification proposed previously [34]. Irregular filaments include strands with rough surface, over-deposited or compressed. The well-defined filaments were further classified as: filaments with equivalent diameter, when the filament has the same diameter as the printing nozzle, and as swollen or stretched filaments, if the diameter is bigger or smaller than the printing nozzle, respectively [34] (see Fig. 3B with marked filament type). Printed PEG-Dop/V^3+^ filaments were irregular at all printing conditions (Fig. 3C, D). The long relaxation time of the network did not allow rearrangement of the crosslinks during shear to adopt the shape imposed by the nozzle. PEG-Dop/Fe^3+^ rendered well-defined strands with diameter between 148 ± 9 and 762 ± 16 at printing pressures between 50 and 500 kPa, whereas PEG-Dop/Al^3+^ formed well-defined strands with diameter between 152 ± 11 and 494 ± 26 within a narrower printing pressure range (25-250 kPa). Printed filaments of PEG-Dop/Al^3+^ at pH 8.4 flowed and merged laterally in few seconds within a significant range of tested conditions (Fig. S6).

The influence of printing pressure and printing speed on the width of the printed filaments is shown in Fig. 3. Increasing printing pressure resulted in wider strands (Fig. 3C, F) which eventually merged. Using 2 mm/s printing speed, strands of PEG-Dop/Al^3+^, PEG-Dop /Fe^3+^ and PEG-Dop /V^3+^ merged when the printing pressure was ≥ 350 kPa, ≥4 50 kPa and ≥ 350 kPa, respectively. Increasing printing speed resulted in thinner filaments, since the amount of deposited material per unit length decreased. At higher printing speeds (5 mm/s for PEG-Dop/Al^3+^ but 10 mm/s for PEG-Dop/Fe^3+^, at 250 kPa printing pressure, Fig. 3D) the printed strands showed discontinuities. PEG-Dop/Al^3+^ and /Fe^3+^ printed filaments at the same speed and pressure showed similar filament width but different edge smoothness (compare Fig. 3C, D, F, G). PEG-Dop/Al^3+^ (pH 8.4), with shorter relaxation time (Fig. S6), rendered significantly wider strand and the filaments merged more easily. If left without secondary crosslinking, all the printed structures fused eventually (Fig. 4B).

**Figure 4.**
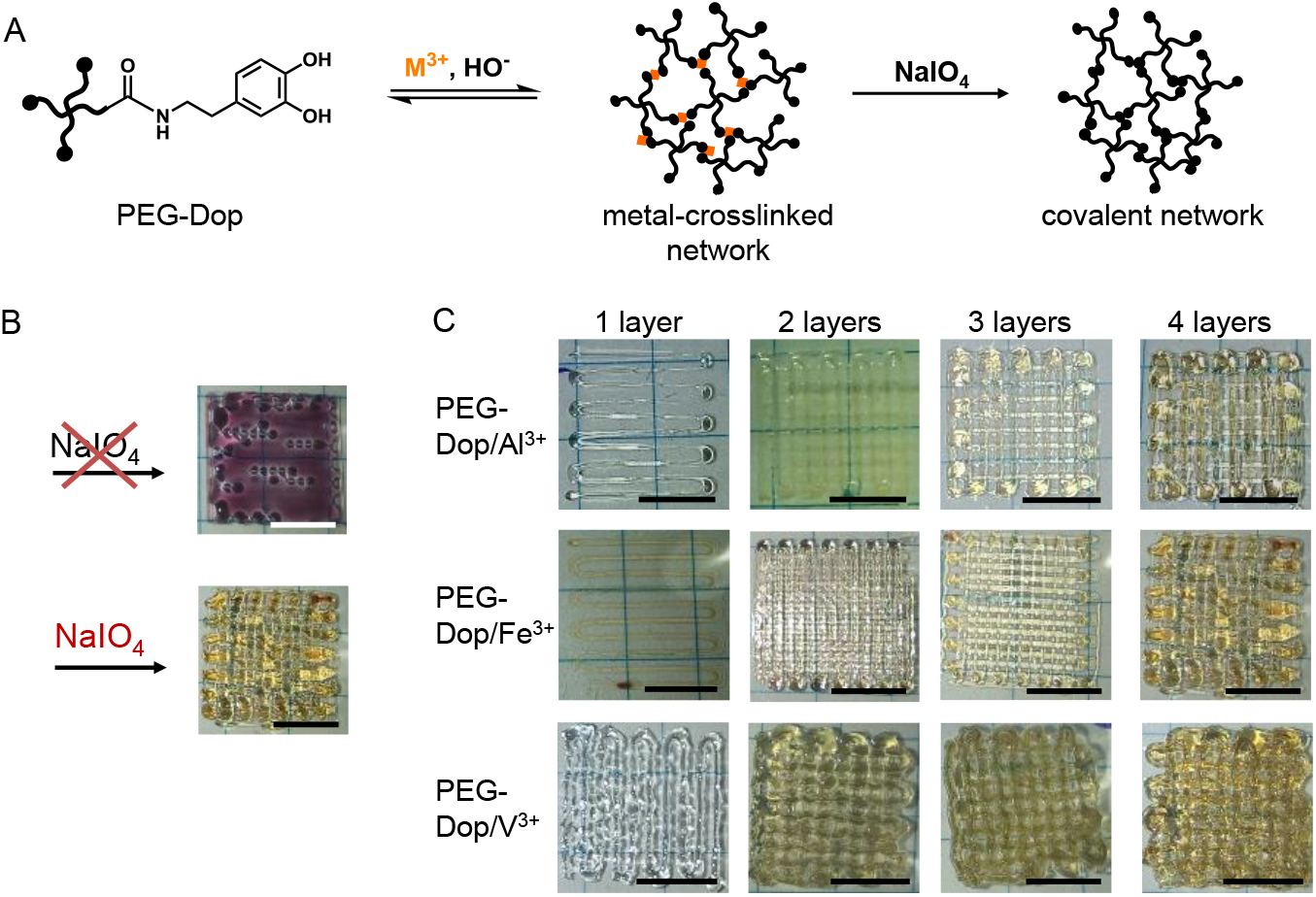
Printing of 3D scaffolds. **A.** Schematic of the secondary covalent crosslinking of the network. **B.** Necessity of introducing covalent crosslinking for permanent shape maintenance, example of 4 layers scaffold printed with PEG-Dop/Fe^3+^ ink (printing conditions: 2 mm/s printing speed at 50 kPa pressure (1^st^ layer) and 100 kPa pressure (2^nd^ - 4^th^ layer)). **C.** Microscopy images of a multilayer construct printed with PEG-Dop/M^3+^ inks. Printing conditions for PEG-Dop/Al^3+^: 3 mm/s printing speed and 85kPa pressure; for a PEG-Dop/Fe^3+^: 2 mm/s printing speed, 50 kPa pressure (1^st^ layer) and 100 kPa pressure (2^nd^ - 4^th^ layer); for PEG-Dop/V^3+^: 2 mm/s printing speed at 200 kPa pressure (1^st^ layer) and 1 mm/s at 250 kPa pressure (2^nd^ - 4^th^ layer). Scale bars: 5 mm.

The thickness of the printed lines can be used as measurement of the accuracy of the printing process [18, 35]. The thinnest printed strand obtained using the nozzle of 260 μm diameter was 148 ± 9, 152 ± 11 and 303 ± 65, for Fe^3+^, Al^3+^ and V^3+^ crosslinked materials, respectively. PEG-Dop/Fe^3+^ ink showed the highest accuracy of the printed structures. Moreover, it also showed the broadest printability range and the smoothest printed structures.

### 3.4 Fabrication of multilayer constructs

The possibility to print 3D structures with PEG-Dop/M^3+^ inks was explored. Due to the reversibility of the metal-catechol crosslinks, printed strands are only temporally stable. In order to increase the mechanical stability of the printed structures and allow 3D stacking of printed strands, a secondary covalently crosslinking step by exposure to mild oxidizing conditions was implemented (Fig. 4). Fig. 4B visualizes how 3D printed layers of filaments of PEG-Dop/Fe^3+^ lose their shape within a few minutes after printing. Addition of an oxidant (sodium periodate) to each layer after printing led to a fast color change and fixation of the printed shape. This is attributed to the self-condensation of the oxidized catechol units to form mechanically stable threads [5] that do not collapse due to own weight. The change in color of the inks containing Al^3+^, Fe^3+^ and V^3+^ from transparent, purple and dark blue respectively, to yellow/orange after treatment with sodium peroxide (Fig. 4B) confirms the presence of quinone oxidized species [13, 36].

1×1 cm^2^ mesh-like constructs consisting of up to 4 layers were successfully printed with PEG-Dop inks crosslinked with Fe^3+^, Al^3+^ and V^3+^ metal ions. Oxidative, covalent crosslinking was performed after deposition of each layer. The higher printing accuracy obtained with PEG-Dop/Fe^3+^ ink in previous experiments with 2D printed structures was also observed in 3D printed scaffolds (Fig. 4C). Additionally, to show the possibility of obtaining bulk 3D scaffolds, pyramid-like constructs consisting of 3 levels, 4-5 layers each, starting from a base area of ~1 cm^2^ were printed (for more details see Supplementary Information, Fig. S7). Covalent crosslinking was performed after each deposited layer. Constructs obtained from PEG-Dop/Fe^3+^ and PEG-Dop/V^3+^ filaments maintained their shape, whereas PEG-Dop/Al^3+^ print deformed. The lower viscosity and poor shape fidelity observed in the 2D printing experiments hinder the construction of 3D structures. These results suggest that inks with higher viscosity allow maintaining the temporary shape for longer times after printing, which is profitable for 3D printing.

### 3.5 Cell viability studies

The viability of cells grown on PEG-Dop networks with metal-coordination crosslinks and with covalent crosslinks was evaluated. Fibroblasts L929 were seeded on the top of PEG-Dop based networks casted in a 96 well-plate. A live/dead assay showed high viability of the cells attached to all tested materials (Fig. 5 and Fig. S8). No statistical differences in viability were observed between cells cultured on the tissue plates and on any of the tested inks. Previous studies have demonstrated toxic effects of Al^3+^ at 2 mM concentration, Fe^3+^ and V^3+^ at 1 mM, for fibroblasts after 48 hours culture [37]. Note that the metal ion concentration in our study was below 1 mM (0.87 mM Me^3+^ concentration, assuming no metal ions removal in washing steps prior to cell seeding). The recommended daily intake for humans of Al, Fe and V ions is ca. 50 mg [38], 6-20 mg [39] and 1.8 mg [40], respectively. This amount would be enough to crosslink large volumes of the presented inks, i.e. 50 ml, 260 ml or 5 ml of the PEG-Dop/Al^3+^, /Fe^3+^ and /V^3+^ inks, respectively.

**Figure. 5.**
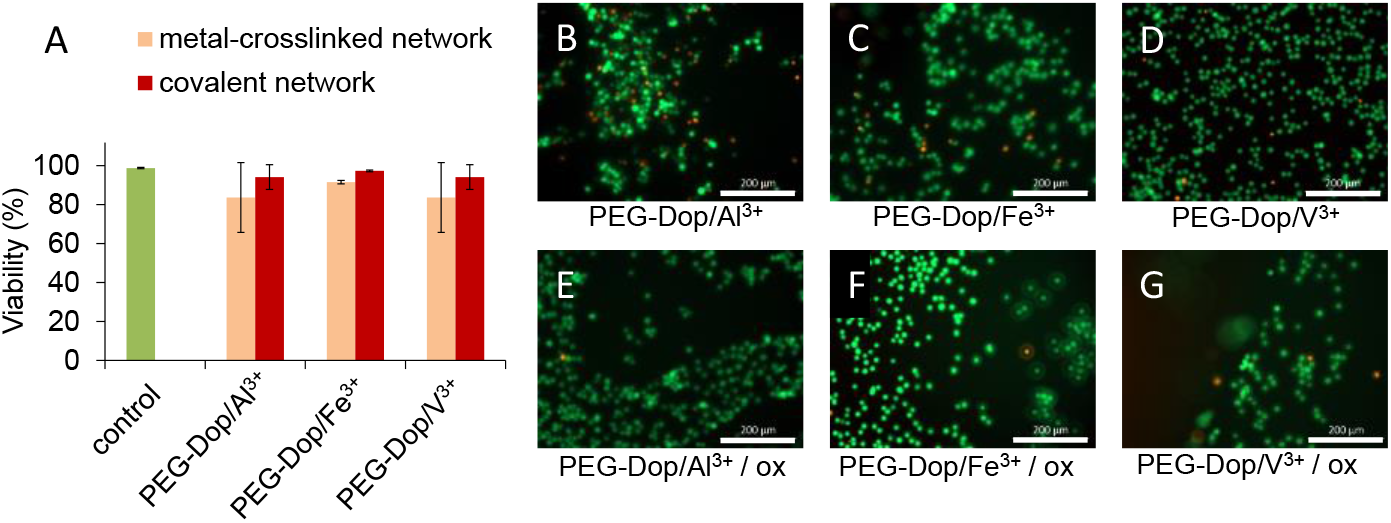
Fibroblasts viability after 1 h culture in direct contact with PEG-Dop based networks. **A.** Viability of L929 cells cultured on top of casted metal-crosslinked PEG-Dop network (orange bars) and covalently crosslinked network by gentle oxidation (red bars), in comparison to cells cultured on plastic (green bar). No significant differences were found. Live/dead staining exemplary images of fibroblasts cultured with metal-crosslinked PEG-Dop networks (**B:** PEG-Dop/Al^3+^, **C:** PEG-Dop/Fe^3+^, **D:** PEG-Dop/V^3+^) and covalent networks crosslinked with oxidant (ox) (**E:** PEG-Dop/Al^3+^, **F:** PEG-Dop/Fe^3+^, **G:** PEG-Dop/V^3+^). Live cells visible in green, dead cells in red. Scale bars: 200 μm.

In the covalently crosslinked gels, we expect that the metal ions are partially depleted from the network due to the consumption of catechol chelating groups during oxidation. The non-bounded cations are expected to be removed, together with remaining oxidant, in the washing step preceding cell seeding on the casted materials. Previous studies showed that addition of 4.5 mg/ml sodium periodate (~20 mM,) to catechol-based hydrogels (final NaIO_4_ concentration ca. 5 mM) during cell encapsulation caused negligible cytotoxicity in vitro and low immunogenicity in vivo. [41] As the NaIO_4_ concentration used in our study was lower than reported for high viability culture [41], no toxicity induced by oxidant was expected even without a washing step. The obtained results indicate that the described printable PEG-Dop/M^3+^ inks are cytocompatible.

## 4. Discussion

The progress of extrusion printing for biomedical applications depends on the development of printable inks that allow ejection at low shear forces to be compatible with living cells, and show fast solidification after extrusion for good shape fidelity and stability of the printed constructs. [4] Systems with reversible bonds combined with a post-printing covalent crosslinking mechanism are suitable systems for this purpose. [2, 4, 10, 11, 42, 43] The reversible bonds lead to printable materials with favorable zero shear viscosities and shear thinning properties for printing. Post-printing covalent crosslinking allows mechanical stabilization of the printed features. Bioinspired catechol-based formulations, which can form reversible metal-coordinated bonds and can autocondensate to form a covalent network at mild oxidative conditions [44], are tested here as an example. Although metal-coordinated systems have been widely applied to many areas of materials science [44], they have only been rarely explored for biofabrication [3, 11, 15]. In this work we have explored ink formulation and printing conditions of PEG-Dop polymers mixed with Al^3+^, Fe^3+^ or V^3+^ ions. This system has been studied previously as dynamic, self-healing network [13, 14].

We identified a suitable formulation for printing consisting of 5% PEG-Dop (10 kDa), 6,67 mM metal cation and 33 mM NaOH final concentrations. The polymer content is remarkably low compared to other printable PEG-based systems (e.g. 10 and 20 % PEG di-methacrylate [45], 25 % p(HPMAm-lactate)-PEG [46], 5-50 % PEG di-acrylate with alginate addition [47], 25 % PEG with 2.5 % alginate addition [48]). A viscoelastic network forms immediately after mixing with the metal ions, and remains stable in time. This is a profitable prerequisite for printing, as the ink can be prepared in advance to the fabrication process and used within few hours without losing its properties.

The properties of the metal coordination bond affect the rheological behavior and printing performance of the PEG-Dop/M^3+^ ink. The tested metal-crosslinked inks (with Al^3+^, Fe^3+^ and V^3+^) allowed extrusion-based printing. Shear forces imposed during printing lead to reversible dissociation of the crosslinks and shear thinning properties, which facilitate extrusion. Differences in printability were observed depending on the properties of the coordination complex used. Al^3+^, Fe^3+^ and V^3+^ complexes with catechol group differ in the coordination degree at a given pH [14] and in the kinetic constant [27], but they show similar thermodynamic stability [25, 49] (Table 2). The **relaxation time** (τ) of the network appeared to be the main parameter influencing the shape and the accuracy of the printed strand. The relaxation time reflects the dynamics and mobility of the network, and the ability to adopt the shape of the nozzle under the shear stress during printing. [4, 30] Well-defined strand deposition was possible with PEG-Dop/Fe^3+^ and PEG-Dop/Al^3+^ networks, with relaxation time below 1 second. PEG-Dop/V^3+^ with relaxation time around 40 s gave irregular structures (see Fig. 3C, D and Fig. 4C), similar to those obtained when printing crosslinked gels [23]. By increasing the speed from 1mm/s to 10 mm/s at a given printed pressure, stretched filaments were obtained for PEG-Dop/Fe^3+^ and PEG-Dop/Al^3+^ networks with relaxation time < 1s. The timescale at which rearrangements occur in the network should match the stretching rate imposed by the moving printing head. The longer τ of PEG-Dop/V^3+^ did not allow for rearrangement in the crosslinking points and lead to breaking of the network during printing in wider range of conditions than for other inks. As a result, discontinuous filaments were obtained more easily. Strands from formulations with relaxation times < 0.1 s, as in PEG-Dop/Al^3+^ ink at pH 8.4, did not retain their shape before post-printing stabilization. In summary, PEG-Dop/M^3+^ networks with relaxation times in the range of one to a few seconds present appropriate balance between easiness of extrusion and good shape fidelity of printed strands which retain the shape until post-printing stabilization by mild oxidative treatment.

The relaxation time of a metal-ligand crosslinked network is expected to be influenced by different parameters characteristic for metal-ligand coordination complex: the coordination degree, the ligand exchange kinetics and the thermodynamic stability constant. A higher coordination degree leads to longer relaxation time, since several ligands interact at the same crosslinking point with the metal cation [13, 14]. Reported studies at pH 8.0 demonstrated that the coordination degree of PEG-Dop/M^3+^ networks decreased in the order of V^3+^>Fe^3+^>Al^3+^. [14] This trend is in agreement with the trend in the relaxation time of the materials observed in our study at pH 8.6-8.9. The coordination degree is expected to decrease with pH, as the ionization state of the catechol unit decreases. In agreement with this, PEG-Dop/Al^3+^ formulation at pH 8.4 showed shorter relaxation times in comparison to the same formulation at pH 8.9. In a first approximation, we expect that the catechol exchange kinetic constant for a catechol/metal ion complex follows the same trend as reported water exchange constants for these complexes [50], i.e. Al^3+^ < Fe^3+^ < V^3+^ (Table 2). Faster kinetics of ligand exchange means more frequent dissociation (and association) of metal-catechol complexes, from which we expected faster network rearrangement [50]. This effect does not explain the longest relaxation time of PEG-Dop/V^3+^. The relation between relaxation times observed for PEG-Dop/M^3+^ networks (PEG-Dop/V^3+^ > PEG-Dop/Fe^3+^ ≈ PEG-Dop/Al^3+^) could be due to the counteracting effects of coordination degree and ligand exchange rate.

The printability differences observed between PEG-Dop/Al^3+^ and PEG-Dop/Fe^3+^ inks cannot be understood based on relaxation time argumentation. We observed that the time needed for stiffness **recovery** after strain-induced breaking of the network was different for the two inks (Fig. 2D). Recovery (self-healing) is possible due to the reversible nature of the metal complexes. [3] The recovery rate is expected to correlate with the rate of ligand exchange. Faster ligand exchange kinetic constants are expected to lead to faster recovery. [50] This trend was observed for all the inks until 70% of the initial stiffness was reached (Fig. 2D and Fig. S2). At higher recovery ratio, deviations from the trend were observed for PEG-Dop/Al^3+^ (pH 8.4) and PEG-Dop/Fe^3+^ (20 kDa). PEG-Dop/Al^3+^ at pH 8.4 recovered faster than expected based on the rate of ligand exchange, presumably due to the lower coordination degree. Faster recovery of PEG-Dop/Fe^3+^ with higher molecular weight could be explained by the lower number of crosslinking points to be restored (1/2 of crosslinks number in 10 kDa PEG network) and the stronger influence of chain entanglements on the final material stiffness. The results suggest that the recovery of the PEG-Dop/M^3+^ networks after strain-induced breaking mostly depends on the healing of bonds and reformation of crosslinks at molecular scale and, therefore, it is dependent on the metal-ligand association rate and not on collective network dynamics (relaxation time), which seems to be more influenced by the metal coordination number. The experimental results suggest that PEG-Dop/M^3+^ with quick recovery time (30% of recovery < 6s) are profitable for shape fidelity. However, too quick recovery may lead to deposition of irregular stands, as in case of PEG-Dop/V^3+^, where full structural rearrangement and therefore well-defined cylindrical shape of strand, was not imposed while extrusion through the printing needle.

We now discuss the relationship between the properties of the coordination bond, the **viscosity** and the printability of the ink. Mathematical modelling of metal coordinated polymeric networks has demonstrated that the viscosity increases exponentially with the coordination degree (as the association of the polymeric chains increases), and decreases with increasing shear rate (shear thinning) as the crosslinks are forced to dissociate. [28] In the tested PEG-Dop/M^3+^ inks, higher values of zero shear viscosity correlated with longer relaxation times and later end of non-Newtonian regime. Shear thinning effect appears above certain critical shear rate *ẏ_c_*, meaning end of non-Newtonian regime, and *ẏ_c_* indicates terminal relaxation time of the polymer. [28] PEG-Dop/V^3+^ had visibly later end of Non-Netwonian regime than PEG-Dop/Fe^3+^ and PEG-Dop/Al^3+^ (Fig. 2C), which is in agreement with much longer relaxation time in comparison to the two other inks. The highest zero shear viscosity was observed for PEG-Dop/V^3+^, the ink with the highest coordination degree. PEG-Dop/Al^3+^ and PEG-Dop/Fe^3+^ showed comparable viscosity in the linear Newtonian regime, with slightly lower values for PEG-Dop/Al^3+^, and even lower for PEG-Dop/Al^3+^ (pH 8.4), indicating that in the system influence of coordination degree on viscosity was observed. The rate of bond dissociation/association is also expected to affect the response of the material to the applied shear. Therefore, viscosity is expected to be influenced by the ligand exchange rate constant: the faster the exchange rate, the higher viscosity. The trend in zero shear viscosity values (PEG-Dop/V^3+^ >> PEG-Dop/Fe^3+^ > PEG-Dop/Al^3+^) follows the order of ligand exchange kinetic constant (V^3+^ > Fe^3+^ >> Al^3+^). To conclude, viscosity influences the printing pressure and the resolution and accuracy of printing. Based on the obtained data, PEG-Dop/M^3+^ with zero shear viscosity in the range of 100 – 1000 Pa·s and showing shear thinning are printable.

Finally, we have observed differences in the elastic response of the inks tested at 10 Hz. PEG-Dop/Fe^3+^ showed higher **elastic modulus** than the PEG-Dop crosslinked with Al^3+^ or V^3+^ (see Table 3). Reported data have related this effect to the higher stability constant (log β) of the complex [13, 36]. However, the difference in the stability constant between the metal-ligand pairs studied here are rather modest [25, 29]. It was shown previously that Fe^3+^ drives catechol oxidation, which leads to introduction of some covalent crosslinks to the network. [14] The fraction of covalent bonds could be accounted for the higher final stiffness of PEG-Dop/Fe^3+^ inks. The elastic behavior in PEG-Dop/M^3+^ networks is dominant at higher frequencies (above 0,025 Hz for PEG-Dop/V^3+^, above ~5 Hz for PEG-Dop/Fe^3+^ and PEG-Dop/Al^3+^), but the behavior of the network at lower frequencies has a predominant influence on printing, i.e. post-printing stability. Therefore, in our system, the elastic modulus is not considered as relevant parameter. In crosslinked gel systems, where G’> G”, sufficient G’ allows to keep the shape of printed form [51] and withstand the weight of consecutive layers. The parameters influencing material rheological properties and recommended values for good printability are collected in Table 4.

**Table 4.** Summary of literature-based relation between catechol-metal complexation chemistry and rheological parameters discussed in the study, and recommended range of these values for good printing based on the study results.

|  | Associated with | Recommended values |
| --- | --- | --- |
| Relaxation time | - **degree of coordination** [13, 14]<br>- **pH** (degree of coordination and competition of HO- for binding sites) [13, 49]<br>- **k** (kinetics of ligand exchange) [30, 50] | ~ 0,2 s |
| Recovery of 30% | - **k** (kinetics of ligand exchange) [50] | 1 s – 6 s |
| Zero shear viscosity | - **degree of coordination** [28]<br>- **pH** (degree of coordination and competition of HO- for binding sites)<br>- **k** (kinetics of ligand exchange) [50] | 100-1000 Pa·s |
| Storage modulus | - **log β** (thermodynamics) [13, 36]<br>- **k** (kinetics of ligand exchange) [49]<br>- polymer **molecular weight** [52] | Not relevant parameter in the presented system |

Catechol-metal ligand crosslinking in PEG-Dop/M^3+^ inks resulted in the shear-thinning and self-healing properties beneficial for printing. However, the reversible character of metal-ligand crosslinking does not provide long-term stability of the printed structures. This is an inherent limitation of printable dynamic networks. [4] However, here the chemical versatility of the catechol group offers an additional advantage: a post-printing treatment under mild oxidative conditions allows covalent stabilization of the structures [5]. In this manner we obtained stable 3D printed structures. Whereas printed filaments of PEG-Dop/Fe^3+^ ink gave the best outcome in terms of well-defined shape and resolution, well-printed 3D scaffolds with the PEG-Dop/V^3+^ were also successfully obtained, indicating that at larger scales the irregular shape of the filaments is no longer relevant. This highlights the importance of understanding the printing process and the ink parameters for the particular application.

## 5. Conclusions

PEG-Dop/M^3+^ based inks show several advantages for 3D extrusion printing. (i) They form reversible networks, with non-covalent crosslinks, that can be stabilized after printing by oxidant-triggered covalent reactions. (ii) The crosslinking units (catechol) can be used for dynamic and covalent crosslinking, and applied to different polymeric systems. There is no need of additional compounds or crosslinking chemistries or UV-triggered reactions, which are the most common ways to introduce post printing covalent crosslinking [42], but can lead to cell damage. (iii) The use of metal ions for crosslinking allows uncomplicated tuning of the material properties by metal exchange, facilitating the adaptation of the system to the requirements of particular printing process (broad range of printing parameters) and application with remarkable flexibility. (iv) We were able to obtain filaments with very good shape fidelity and resolution down to 150 μm. Thinner filaments might be obtained using printing nozzles with smaller diameters. (v) The PEG-Dop/M^3+^ inks require low polymer content (final: 5%), following one of the most stringent criteria in bioink design [2].

The development of functional bioinks is a challenging task. Our results regarding the influence of crosslinking chemistry and reversible bond parameters on the rheological properties and printability of the ink will guide material design in other reversible systems. We defined relaxation time and recovery time as main parameters indicating printability of ink with reversible bonds.

## Acknowledgements

The authors thank Prof. Santiago J. Garcia from the Faculty of Aerospace Engineering at the TU Delft, The Netherlands and Prof. Christian Wagner from Experimental Physics, Saarland University, Saarbrücken, Germany, for the stimulating and fruitful discussions on rheological data. J.F. acknowledges the financial support from China Scholarship Council. J.I.P., M.V. and A.d.C. acknowledge financial support from the EU within BioSmartTrainee, the Marie Skłodowska-Curie Innovative Training School, No. 642861.

## References

[1] A. Ribeiro, M.M. Blokzijl, R. Levato, C.W. Visser, M. Castilho, W.E. Hennink, T. Vermonden, J. Malda, Assessing bioink shape fidelity to aid material development in 3D bioprinting, Biofabrication 10(1) (2018) 014102.

[2] M. Jos, V. Jetze, M.F. P., J. Tomasz, H.W. E., D.W.J. A., G. Jürgen, H.D. W., 25th Anniversary Article: Engineering Hydrogels for Biofabrication, Advanced Materials 25(36) (2013) 5011–5028.

[3] S.Y. Zheng, H. Ding, J. Qian, J. Yin, Z.L. Wu, Y. Song, Q. Zheng, Metal-Coordination Complexes Mediated Physical Hydrogels with High Toughness, Stick–Slip Tearing Behavior, and Good Processability, Macromolecules 49(24) (2016) 9637–9646.

[4] T. Jungst, W. Smolan, K. Schacht, T. Scheibel, J. Groll, Strategies and Molecular Design Criteria for 3D Printable Hydrogels, Chemical Reviews 116(3) (2016) 1496–1539.

[5] J.I. Paez, O. Ustahüseyin, C. Serrano, X.-A. Ton, Z. Shafiq, G.K. Auernhammer, M. d’Ischia, A. del Campo, Gauging and Tuning Cross-Linking Kinetics of Catechol-PEG Adhesives via Catecholamine Functionalization, Biomacromolecules 16(12) (2015) 3811–3818.

[6] L.J. Macdougall, M.M. Pérez-Madrigal, M.C. Arno, A.P. Dove, Nonswelling Thiol–Yne Cross-Linked Hydrogel Materials as Cytocompatible Soft Tissue Scaffolds, Biomacromolecules 19(5) (2018) 1378–1388.

[7] T. Atabak Ghanizadeh, A.H. Miguel, R.L. Nicholas, S. Wenmiao, Three-dimensional bioprinting of complex cell laden alginate hydrogel structures, Biofabrication 7(4) (2015) 045012.

[8] L.L. Wang, C.B. Highley, Y.-C. Yeh, J.H. Galarraga, S. Uman, J.A. Burdick, Three-dimensional extrusion bioprinting of single- and double-network hydrogels containing dynamic covalent crosslinks, Journal of Biomedical Materials Research Part A 106(4) (2018) 865–875.

[9] C.B. Highley, C.B. Rodell, J.A. Burdick, Direct 3D Printing of Shear-Thinning Hydrogels into Self-Healing Hydrogels, Advanced Materials 27(34) (2015) 5075–5079.

[10] M. Shin, J.H. Galarraga, M.Y. Kwon, H. Lee, J.A. Burdick, Gallol-derived ECM-mimetic adhesive bioinks exhibiting temporal shear-thinning and stabilization behavior, Acta Biomaterialia (2018).

[11] L. Shi, H. Carstensen, K. Hölzl, M. Lunzer, H. Li, J. Hilborn, A. Ovsianikov, D.A. Ossipov, Dynamic Coordination Chemistry Enables Free Directional Printing of Biopolymer Hydrogel, Chemistry of Materials 29(14) (2017) 5816–5823.

[12] M. Krogsgaard, M.A. Behrens, J.S. Pedersen, H. Birkedal, Self-Healing Mussel-Inspired Multi-pH-Responsive Hydrogels, Biomacromolecules 14(2) (2013) 297–301.

[13] N. Holten-Andersen, M.J. Harrington, H. Birkedal, B.P. Lee, P.B. Messersmith, K.Y. Lee, J.H. Waite, pH-induced metal-ligand cross-links inspired by mussel yield self-healing polymer networks with near-covalent elastic moduli, Proceedings of the National Academy of Sciences 108(7) (2011) 2651–5.

[14] N. Holten-Andersen, A. Jaishankar, M.J. Harrington, D.E. Fullenkamp, G. DiMarco, L.H. He, G.H. McKinley, P.B. Messersmith, K.Y.C. Leei, Metal-coordination: using one of nature’s tricks to control soft material mechanics, Journal of Materials Chemistry B 2(17) (2014) 2467–2472.

[15] X. Li, H. Wang, D. Li, S. Long, G. Zhang, Z. Wu, Dual Ionically Cross-linked Double-Network Hydrogels with High Strength, Toughness, Swelling Resistance, and Improved 3D Printing Processability, ACS Applied Materials & Interfaces 10(37) (2018) 31198–31207.

[16] L. Shi, Y. Zhao, Q. Xie, C. Fan, J. Hilborn, J. Dai, D.A. Ossipov, Moldable Hyaluronan Hydrogel Enabled by Dynamic Metal–Bisphosphonate Coordination Chemistry for Wound Healing, Advanced Healthcare Materials 7(5) (2018) 1700973.

[17] J. Feng, X.-A. Ton, S. Zhao, J. Paez, A. del Campo, Mechanically Reinforced Catechol-Containing Hydrogels with Improved Tissue Gluing Performance, Biomimetics 2(4) (2017) 23.

[18] H. Li, S. Liu, L. Lin, Rheological study on 3D printability of alginate hydrogel and effect of graphene oxide, International Journal of Bioprinting 2(2) (2016) 54–66.

[19] H. Li, Y.J. Tan, S. Liu, L. Li, Three-Dimensional Bioprinting of Oppositely Charged Hydrogels with Super Strong Interface Bonding, ACS Applied Materials & Interfaces 10(13) (2018) 11164–11174.

[20] H.J. Li, Y.J. Tan, K.F. Leong, L. Li, 3D Bioprinting of Highly Thixotropic Alginate/Methylcellulose Hydrogel with Strong Interface Bonding, ACS Applied Materials & Interfaces 9(23) (2017) 20086–20097.

[21] N. Diamantides, L. Wang, T. Pruiksma, J. Siemiatkoski, C. Dugopolski, S. Shortkroff, S. Kennedy, L.J. Bonassar, Correlating rheological properties and printability of collagen bioinks: the effects of riboflavin photocrosslinking and pH, Biofabrication 9(3) (2017) 034102.

[22] N. Paxton, W. Smolan, T. Bock, F. Melchels, J. Groll, T. Jungst, Proposal to assess printability of bioinks for extrusion-based bioprinting and evaluation of rheological properties governing bioprintability, Biofabrication 9(4) (2017) 044107.

[23] L. Ouyang, R. Yao, Y. Zhao, W. Sun, Effect of bioink properties on printability and cell viability for 3D bioplotting of embryonic stem cells, Biofabrication 8(3) (2016) 035020.

[24] L. Garcia-Fernandez, J. Cui, C. Serrano, Z. Shafiq, R.A. Gropeanu, V.S. Miguel, J.I. Ramos, M. Wang, G.K. Auernhammer, S. Ritz, A.A. Golriz, R. Berger, M. Wagner, A. del Campo, Antibacterial strategies from the sea: polymer-bound cl-catechols for prevention of biofilm formation, Advanced Materials 25(4) (2013) 529–33.

[25] V.M. Nurchi, T. Pivetta, J.I. Lachowicz, G. Crisponi, Effect of substituents on complex stability aimed at designing new iron(III) and aluminum(III) chelators, Journal of Inorganic Biochemistry 103(2) (2009) 227–236.

[26] M.J. Sever, J.J. Wilker, Absorption spectroscopy and binding constants for first-row transition metal complexes of a DOPA-containing peptide, Dalton Transactions (6) (2006) 813–822.

[27] L. Helm, A. Merbach, Applications of advanced experimental techniques: high pressure NMR and computer simulations., Journal of the Chemical Society, Dalton Transactions 5 (2002) 633–641.

[28] K.P. Santo, A. Vishnyakov, R. Kumar, A.V. Neimark, Elucidating the Effects of Metal Complexation on Morphological and Rheological Properties of Polymer Solutions by a Dissipative Particle Dynamics Model, Macromolecules 51(14) (2018) 4987–5000.

[29] M.S. Menyo, C.J. Hawker, J.H. Waite, Rate-Dependent Stiffness and Recovery in Interpenetrating Network Hydrogels through Sacrificial Metal Coordination Bonds, ACS Macro Letters 4(11) (2015) 1200–1204.

[30] R.K. Bose, N. Hohlbein, S.J. Garcia, A.M. Schmidt, S. van der Zwaag, Connecting supramolecular bond lifetime and network mobility for scratch healing in poly(butyl acrylate) ionomers containing sodium, zinc and cobalt, Physical Chemistry Chemical Physics 17(3) (2015) 1697–704.

[31] J. Wei, J. Wang, S. Su, S. Wang, J. Qiu, Z. Zhang, G. Christopher, F. Ning, W. Cong, 3D printing of an extremely tough hydrogel, RSC Advances 5(99) (2015) 81324–81329.

[32] T. Hoare, D. Zurakowski, R. Langer, D.S. Kohane, Rheological blends for drug delivery. I. Characterization in vitro, Journal of Biomedical Materials Research Part A 92A(2) (2010) 575–585.

[33] R. Suntornnond, E.Y.S. Tan, J. An, C.K. Chua, A Mathematical Model on the Resolution of Extrusion Bioprinting for the Development of New Bioinks, Materials, 2016.

[34] Y. Jin, W. Chai, Y. Huang, Printability study of hydrogel solution extrusion in nanoclay yield-stress bath during printing-then-gelation biofabrication, Materials Science and Engineering: C 80 (2017) 313–325.

[35] B. Webb, B.J. Doyle, Parameter optimization for 3D bioprinting of hydrogels, Bioprinting 8 (2017) 8–12.

[36] M. Krogsgaard, M.R. Hansen, H. Birkedal, Metals & polymers in the mix: fine-tuning the mechanical properties & color of self-healing mussel-inspired hydrogels Journal of Materials Chemistry B 2(47) (2014) 8292–8297.

[37] N.J. Hallab, S. Anderson, M. Caicedo, A. Brasher, K. Mikecz, J.J. Jacobs, Effects of soluble metals on human peri-implant cells, Journal of Biomedical Materials Research Part A 74A(1) (2005) 124–140.

[38] D. Krewski, R.A. Yokel, E. Nieboer, D. Borchelt, J. Cohen, J. Harry, S. Kacew, J. Lindsay, A.M. Mahfouz, V. Rondeau, Human health risk assessment for aluminium, aluminium oxide, and aluminium hydroxide, Journal of Toxicology and Environmental Health Part B: Critical Reviews 10 Suppl 1 (2007) 1–269.

[39] K. Prasad, O. Bazaka, M. Chua, M. Rochford, L. Fedrick, J. Spoor, R. Symes, M. Tieppo, C. Collins, A. Cao, D. Markwell, K.K. Ostrikov, K. Bazaka, Metallic Biomaterials: Current Challenges and Opportunities, Materials 10 (2017) 884.

[40] Opinion of the Scientific Panel on Dietetic products, nutrition and allergies [NDA] related to the Tolerable Upper Intake Level of Vanadium, EFSA Journal 2(3) (2004) 33.

[41] C. Lee, J. Shin, J.S. Lee, E. Byun, J.H. Ryu, S.H. Um, D.-I. Kim, H. Lee, S.-W. Cho, Bioinspired, Calcium-Free Alginate Hydrogels with Tunable Physical and Mechanical Properties and Improved Biocompatibility, Biomacromolecules 14(6) (2013) 2004–2013.

[42] D. Petta, D.W. Grijpma, M. Alini, D. Eglin, M. D’Este, Three-Dimensional Printing of a Tyramine Hyaluronan Derivative with Double Gelation Mechanism for Independent Tuning of Shear Thinning and Postprinting Curing, ACS Biomaterials Science & Engineering 4(8) (2018) 3088–3098.

[43] L. Ouyang, C.B. Highley, C.B. Rodell, W. Sun, J.A. Burdick, 3D Printing of Shear-Thinning Hyaluronic Acid Hydrogels with Secondary Cross-Linking, ACS Biomaterials Science & Engineering 2(10) (2016) 1743–1751.

[44] B. Wang, Y. S. Jeon, H.S. Park, Y. J. Kim, J.-H. Kim, Mussel-mimetic self-healing polyaspartamide derivative gel via boron-catechol interactions, Express Polymer Letters 9 (2015) 799–808.

[45] X. Cui, K. Breitenkamp, M.G. Finn, M. Lotz, D.D. D’Lima, Direct Human Cartilage Repair Using Three-Dimensional Bioprinting Technology, Tissue Engineering Part A 18(11–12) (2012) 1304–1312.

[46] R. Censi, W. Schuurman, J. Malda, G. di Dato, P.E. Burgisser, W.J.A. Dhert, C.F. van Nostrum, P. di Martino, T. Vermonden, W.E. Hennink, A Printable Photopolymerizable Thermosensitive p(HPMAm-lactate)-PEG Hydrogel for Tissue Engineering, Advanced Functional Materials 21(10) (2011) 1833–1842.

[47] L.A. Hockaday, K.H. Kang, N.W. Colangelo, P.Y.C. Cheung, B. Duan, E. Malone, J. Wu, L.N. Girardi, L.J. Bonassar, H. Lipson, C.C. Chu, J.T. Butcher, Rapid 3D printing of anatomically accurate and mechanically heterogeneous aortic valve hydrogel scaffolds, Biofabrication 4(3) (2012) 035005.

[48] S. Hong, D. Sycks, H.F. Chan, S. Lin, G.P. Lopez, F. Guilak, K.W. Leong, X. Zhao, 3D Printing of Highly Stretchable and Tough Hydrogels into Complex, Cellularized Structures, Advanced Materials 27(27) (2015) 4035–4040.

[49] M.S. Menyo, C.J. Hawker, J.H. Waite, Versatile tuning of supramolecular hydrogels through metal complexation of oxidation-resistant catechol-inspired ligands, Soft Matter 9(43) (2013).

[50] D.E. Fullenkamp, L. He, D.G. Barrett, W.R. Burghardt, P.B. Messersmith, Mussel-inspired histidine-based transient network metal coordination hydrogels, Macromolecules 46(3) (2013) 1167–1174.

[51] J.H.Y. Chung, S. Naficy, Z.L. Yue, R. Kapsa, A. Quigley, S.E. Moulton, G.G. Wallace, Bio-ink properties and printability for extrusion printing living cells, Biomaterials Science 1(7) (2013) 763–773.

[52] S.C. Grindy, M. Lenz, N. Holten-Andersen, Engineering Elasticity and Relaxation Time in Metal-Coordinate Cross-Linked Hydrogels, Macromolecules 49(21) (2016) 8306–8312.

