## Supplementary Information for "Printability study of metal ion crosslinked PEG-catechol based inks"

1) <u>Defining ink compositions used for the study</u>

**Table S1.** Tested compositions for printability. PEG-Dop, $Fe^{3+}$ ion and NaOH concentration values in the table refer to the solutions before mixing. The formulation selected for further studies is indicated in yellow.

| initial [PEG-Dop] | initial [$Fe^{3+}$] | cat:$Fe^{3+}$ ratio | initial [NaOH] | network formation |
|---|---|---|---|---|
| 20 % | 80 mM | (3:1) | 1 mM | no |
| | | | 5 mM | no |
| | | | 10 mM | no |
| | | | 50 mM | yes, inhomogeneous |
| | | | 100 mM | yes, inhomogeneous |
| | | | 250 mM* | yes, homogeneous |
| | | | 1 M | yes, inhomogeneous |
| 10 % | 40mM | (3:1) | 50 mM | yes, inhomogeneous |
| | | | 100 mM * | yes, homogeneous |
| | | | 250 mM | yes, homogeneous |
| | | | 1 M | yes, inhomogeneous |
| 5 % | 20mM | (3:1) | 50 mM * | yes, homogeneous |
| | | | 100 mM | yes, inhomogeneous |
| | | | 250 mM | yes, inhomogeneous |
| | | | 1 M | yes, inhomogeneous |

* Samples that led to homogenous network formation at the lowest tested NaOH content

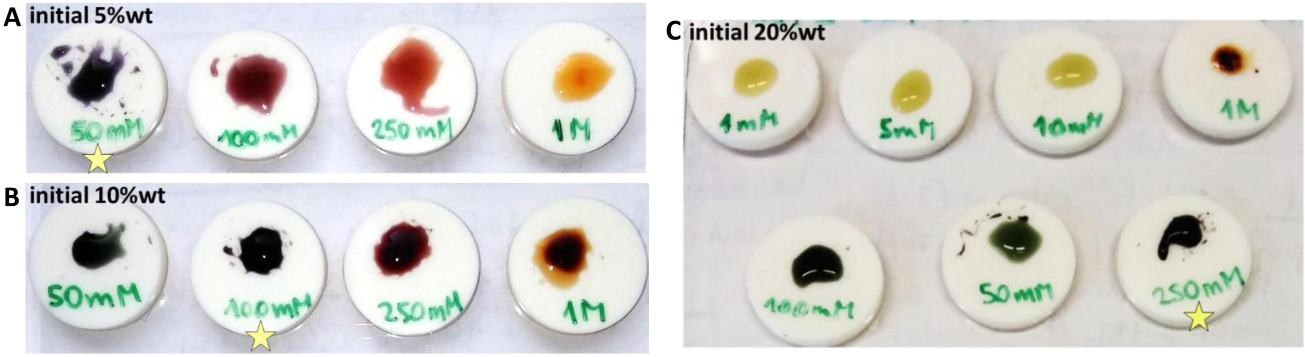


**Figure S1**. Images of PEG-Dop networks crosslinked with $Fe^{3+}$. Initial PEG-Dop content: **A** – 5 %, **B** – 10 %, **C** – 20%. Initial NaOH concentration in each composition is indicated below the mixtures. The best conditions for formation of a homogeneous network (at lowest possible pH in view of future biomedical applications) are indicated with the star. Based on the preliminary extrusion test with 1 ml syringe, network with 5% initial polymer concentration was assessed as too low viscosity, whereas 20% caused clogging of the syringe. 10% was chosen for final studies. To keep comparable conditions, same concentrations of the compounds were used for obtaining inks crosslinked with $Al^{3+}$ and $V^{3+}$ ions.

2) Rheological characterization of tested inks

**Table S2.** Relaxation time and storage modulus of all tested inks.

| Ink | Relaxation time ($\tau$) (s) | Storage modulus ($G'$) (Pa)* |
|---|---|---|
| PEG-Dop/$Al^{3+}$ | 0,23 ± 0,02 | 4020 ± 973 |
| PEG-Dop/$Al^{3+}$ (pH 8.4) | 0,12 ± 0,01 | 2045 ± 463 |
| PEG-Dop/$Fe^{3+}$ | 0,20 ± 0,06 | 7233 ± 1406 |
| PEG-Dop/$Fe^{3+}$ (20 kDa) | 0,18 ± 0,02 | 4777 ± 1179 |
| PEG-Dop/$V^{3+}$ | 44 ± 19 | 2732 ± 1162 |

* Measured at 10 Hz frequency and 1% strain

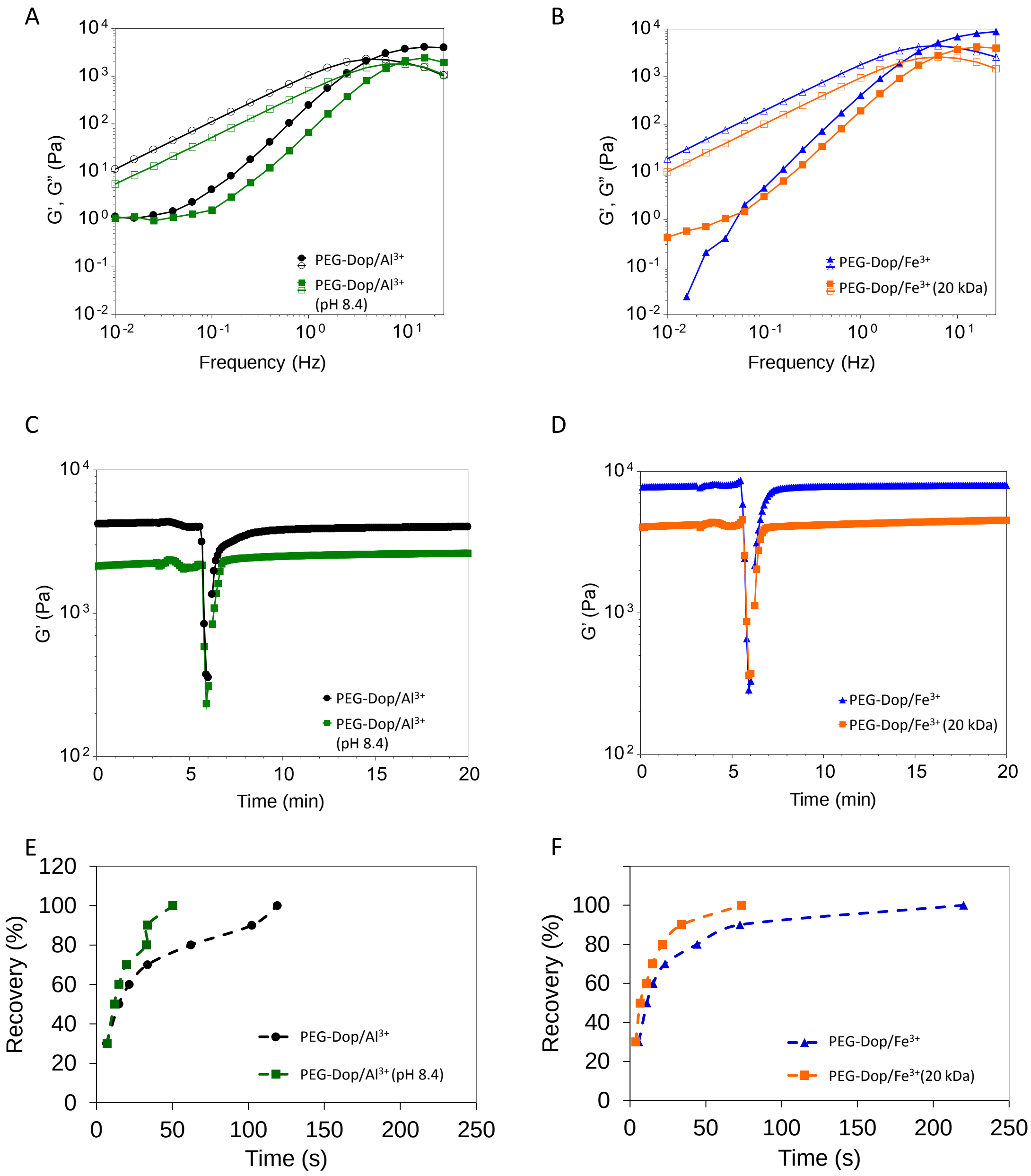


**Figure S2.** Rheological curves of PEG-Dop networks with metal complexation crosslinking; comparison between PEG-Dop/$Al^{3+}$ and PEG-Dop/$Al^{3+}$ (pH 8.4), PEG-Dop/$Fe^{3+}$ and PEG-Dop/$Fe^{3+}$ (20 kDa). The storage (full symbols) and loss (open symbols) modulus, G' and G", recorded as a function of frequency at 1% strain for: PEG-Dop/$Al^{3+}$ and PEG-Dop/$Al^{3+}$ (pH 8.4) (**A**), PEG-Dop/$Fe^{3+}$ and PEG-Dop/$Fe^{3+}$ (20 kDa) (**B**). Time sweep for: PEG-Dop/$Al^{3+}$ and PEG-Dop/$Al^{3+}$ (pH 8.4) (**C**), and PEG-Dop/$Fe^{3+}$ and PEG-Dop/$Fe^{3+}$ (20 kDa) (**D**), measured at 10 Hz frequency and 1% strain, followed by initiated after 3 min logarithmic strain increase from 0.01 to 1000%, leading to network breakage at ~5.5 min. After 6 minutes network recovery was recorded in the consequent time sweep measurement at 10 Hz frequency and 1% strain. Average time necessary for the recovery of: PEG-Dop/$Al^{3+}$ and PEG-Dop/$Al^{3+}$ (pH 8.4) (**E**), and PEG-Dop/$Fe^{3+}$ and PEG-Dop/$Fe^{3+}$ (20 kDa) (**F**) after network breaking by the applied strain increasing from 0.01 to 1000%. The lines serve as a guide for the eye.

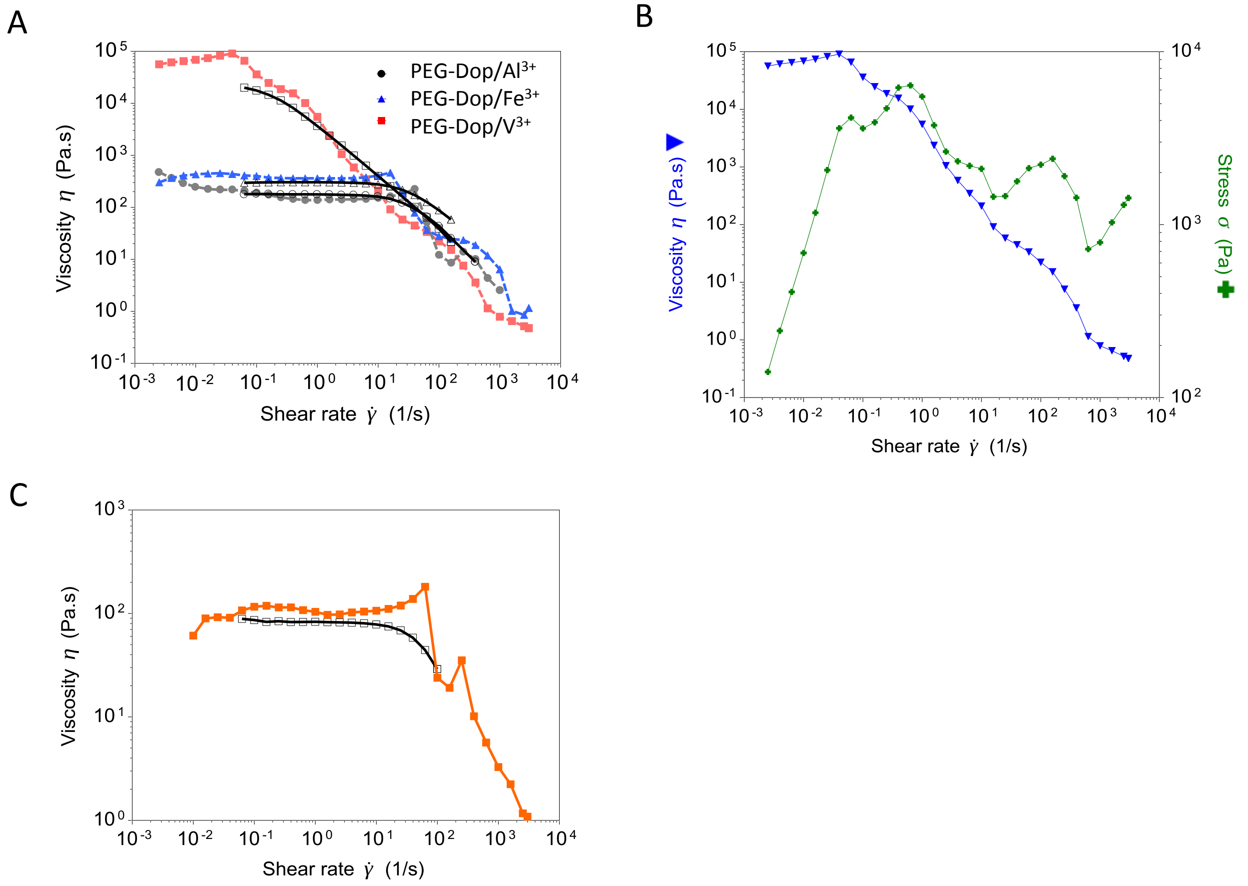


**Figure S3.** Results of the viscosity measurements at increasing share rate, performed in the rotational flow sweep experiment. **A.** Comparison between data obtained directly in the rotational flow sweep experiment (full symbols) and results obtained from application of Cox-Merz rule to oscillatory data (open symbols), for PEG-Dop/$Al^{3+}$ (circles), PEG-Dop/$Fe^{3+}$ (tirangles), PEG-Dop/$V^{3+}$ (squares) inks. The data are corresponding well. **B.** Representative viscosity measurement for PEG-Dop/$V^{3+}$ ink (triangles) with plotted stress values (crosses) obtained at increasing shear rate, indicating sample edge failure, confirmed by macroscopic observation. Similar stress dependency was observed for all other tested samples. **C.** Comparison between data obtained directly in the rotational flow sweep experiment (full symbols) and results obtained from application of Cox-Merz rule to oscillatory data (open symbols) for PEG-Dop/$Al^{3+}$ (pH 8.4) ink. Note clearly visible differences in the shear rate in the range of $10^2 \cdot 1/s$. For all other tested inks this discrepancy did not occur.

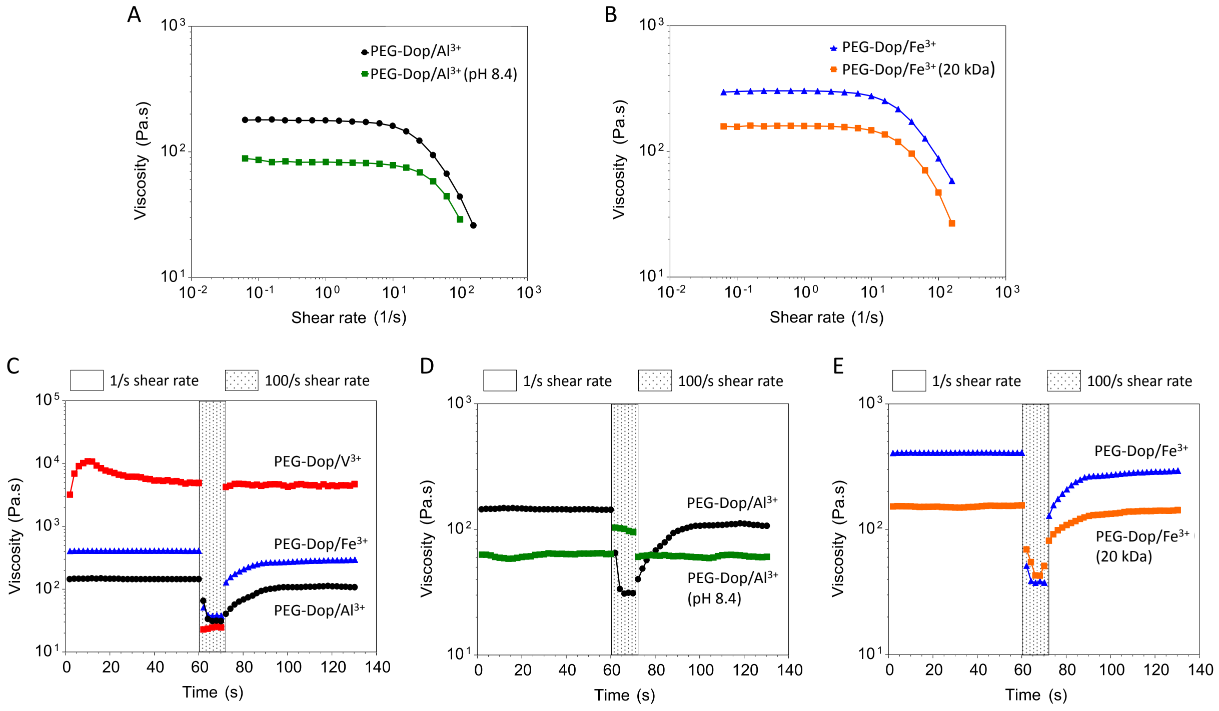


**Figure S4.** Shear thinning and viscosity recovery properties. Viscosity of the inks as a function of shear rate obtained from Cox-Merz rule applied to oscillatory data for: PEG-Dop/$Al^{3+}$ and PEG-Dop/$Al^{3+}$ (pH 8.4) (**A**), PEG-Dop/$Fe^{3+}$ and PEG-Dop/$Fe^{3+}$ (20 kDa) (**B**). Recovery of viscosity measured for: PEG-Dop/$Al^{3+}$, PEG-Dop/$Fe^{3+}$ and PEG-Dop/$V^{3+}$ inks (**C**); PEG-Dop/$Al^{3+}$ and PEG-Dop/$Al^{3+}$ (pH 8.4) (**D**); PEG-Dop/$Fe^{3+}$ and PEG-Dop/$Fe^{3+}$ (20 kDa) (**E**). To monitor recovery of viscosity a step shear rate test was performed on the freshly prepared samples. First, the sample viscosity was monitored for 60 s at a constant (low) shear rate of 1/s, followed by (high) shear rate of 100 · 1/s applied for 10 s. Viscosity recovery was followed afterwards for 60 s at initial shear rate of 1/s. Note the increase in viscosity for PEG-Dop/$Al^{3+}$ (pH 8.4) at 100 · 1/s. This effect can be explained by visible increase of viscosity in flow sweep measurement (Fig.S3C) at the shear rates ~ 100 · 1/s, before revealed shear thinning.

3) Determination of maximum shear rate and apparent viscosity while printing

For a non-Newtonian fluid in a capillary viscometer, shear stress can be defined as [1]:

$$\tau = m(\dot{\gamma})^n \qquad \textit{(Equation 1)}$$

where:

$\tau$ – shear stress on the wall of the capillary;
$\dot{\gamma}$ – shear rate on the wall of the capillary;
*n* – power law index;
*m* – power law consistency coefficient;

The viscosity can be respectively described as:

$$\eta = m(\dot{\gamma})^{n-1} \qquad \textit{(Equation 2)}$$

where:

$\eta$ – apparent viscosity;

The maximum shear rate (shear rate on the wall of the capillary/nozzle) will follow the relation [2]:

$$\dot{\gamma} = \frac{3n+1}{4n} \cdot \frac{4\dot{Q}}{\pi R^3} = \frac{3n+1}{4n} \cdot \frac{32\dot{Q}}{\pi D^3} \qquad \textit{(Equation 3)}$$

where:

$\eta$ – viscosity;
$\dot{Q}$ – volumetric flow rate;
R – nozzle radius;
D – nozzle diameter;

For a strand printed continuously, with homogeneous geometry, the flow rate can be found from:

$$\dot{Q} = \frac{\pi d^2}{4}\dot{v} \qquad \textit{(Equation 4)}$$

where:

$\dot{v}$ – printing head moving speed;
d – printed filament diameter;

Based on *Equitation 3* and *Equation 4*, one can obtain:

$$\dot{\gamma} = \frac{3n+1}{4n} \cdot \frac{8d^2}{D^3}\dot{v} \qquad \textit{(Equation 5)}$$

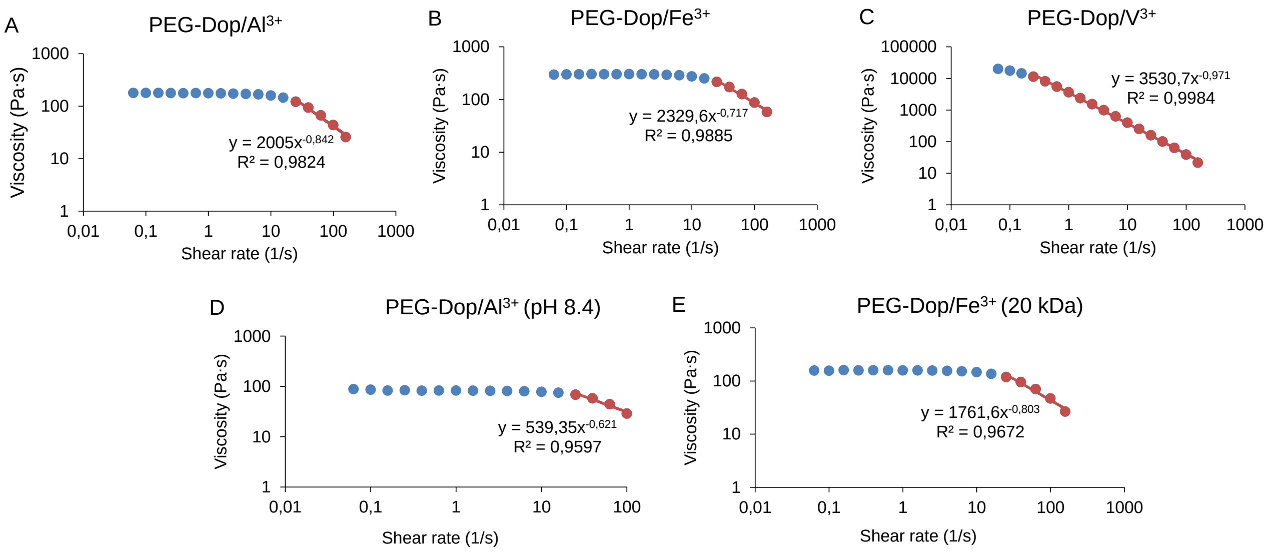


**Figure S5.** Viscosity as a function of a shear rate based on the Cox-Merz rule applied to the oscillatory measurements of frequency sweep, and power-law fit for the shear thinning region (indicated in red), for different samples: **A**. PEG-Dop/$V^{3+}$, **B**. PEG-Dop/$Fe^{3+}$, **C**. PEG-Dop/$Al^{3+}$, **D**. PEG-Dop/$Al^{3+}$ (pH 8.4), **E**. PEG-Dop/$Fe^{3+}$ (20 kDa).

**Table S3.** Theoretical maximum shear rate experienced by the ink while printing, calculated according to *Equation 5*, and corresponding viscosity.

| Material | Power law index (*n*) | Head moving speed (mm/s) | Maximum shear rate (1/s) | Viscosity (Pa·s) |
|---|---|---|---|---|
| PEG-Dop/$V^{3+}$ | 0,029 | 1 | 72 [b] | 55 |
| | | 2 | 144 [b] | 28 |
| | | 5 | 360 [b] | 12 |
| | | 10 | 721 [b] | 6 |
| PEG-Dop/$Fe^{3+}$ | 0,283 | 1 | 8 [a] | 280 |
| | | 2 | 25 [b] | 231 |
| | | 5 | 63 [b] | 120 |
| | | 10 | 126 [b] | 73 |
| PEG-Dop/$Fe^{3+}$ (20 kDa) | 0,197 | 1 | 8 [a] | 150 |
| | | 2 | 31 [b] | 112 |
| | | 5 | 78 [b] | 53 |
| | | 10 | 155 [b] | 31 |
| PEG-Dop/$Al^{3+}$ | 0,158 | 1 | 8 [a] | 164 |
| | | 2 | 36 [b] | 98 |
| | | 5 | 90 [b] | 45 |
| | | 10 | 179 [b] | 25 |
| PEG-Dop/$Al^{3+}$ (pH 8.4) | 0,379 | 1 | 8 [a] | 79 |
| | | 2 | 15 [a] | 75 |
| | | 5 | 54 [b] | 45 |
| | | 10 | 108 [b] | 29 |

a - Newtonian range, b – non-Newtonian range

4) Quantification of printability

**Table S4.** Strand width obtained at all attempted printed conditions (pressure and speed) for all tested inks.

| Press (kPa) | Speed (mm/s) | PEG-Dop/$V^{3+}$ | PEG-Dop/$Fe^{3+}$ | PEG-Dop/$Al^{3+}$ | PEG-Dop/$Al^{3+}$ (pH 8.4) | PEG-Dop/$Fe^{3+}$ (20 kDa) |
|---|---|---|---|---|---|---|
| 15 | 1 | N/A [a] | 78±65 [a] | 82±49 [a] | **200±8** | 46±13 [a] |
| | 2 | - | 49±42 [a] | 50±39 [a] | 128±95 [a] | 60±48 [a] |
| | 5 | - | N/A [a] | N/A [a] | 71±50 [a] | 32±28 [a] |
| | 10 | - | N/A [a] | N/A [a] | 25±27 [a] | N/A [a] |
| 25 | 1 | N/A [a] | 133±65 [a] | **152±11** | **306±9** | **202±8** |
| | 2 | - | 91±51 [a] | 102±29 [a] | **210±9** | 90±57 [a] |
| | 5 | - | N/A [a] | N/A [a] | 103±76 [a] | 64±31 [a] |
| | 10 | - | N/A [a] | N/A [a] | 73±63 [a] | 21±29 [a] |
| 35 | 1 | N/A [a] | 161±63 [a] | **207±28** | **345±23** | **251±11** |
| | 2 | - | 73±49 [a] | 142±20 [a] | **232±12** | 181±6 [a] |
| | 5 | - | 56±29 [a] | N/A [a] | 91±20 [a] | 90±51 [a] |
| | 10 | - | N/A [a] | N/A [a] | N/A [a] | 57±55 [a] |
| 50 | 1 | N/A [a] | **222±8** | 255±21 [c] | **433±25** | 318±10 [c] |
| | 2 | - | **148±9** | 157±28 [c] | **296±10** | **227±9** |
| | 5 | - | N/A [a] | N/A [a] | **184±12** | 142±36 [a] |
| | 10 | - | N/A [a] | N/A [a] | 109±65 [a] | 55±62 [a] |
| 75 | 1 | N/A [a] | **278±8** | **310±19** | N/A [b] | **397±15** |
| | 2 | - | **189±9** | 206±29 [c] | **349±13** | **265±11** |
| | 5 | - | 88±106 [a] | N/A [a] | **199±11** | 167±8 [c] |
| | 10 | - | N/A [a] | N/A [a] | N/A [a] | 74±64 [a] |
| 85 | 1 | N/A [a] | **296±9** | **308±22** | N/A [b] | **399±12** |
| | 2 | - | **191±13** | **201±25** | **349±13** | **286±5** |
| | 5 | - | 95±82 [a] | N/A [a] | **215±16** | 168±13 [c] |
| | 10 | - | N/A [a] | N/A [a] | N/A [a] | 99±73 [a] |
| 100 | 1 | - | **308±13** | **350±37** | N/A [b] | **457±16** |
| | 2 | - | **198±16** | 202±38 [c] | **392±10** | **301±13** |
| | 5 | - | 123±105 [a] | N/A [a] | **217±11** | **184±16** |
| | 10 | - | N/A [a] | N/A [a] | 131±79 [a] | 96±125 [a] |
| 120 | 1 | N/A [a] | **330±11** | **346±16** | - | **506±16** |
| | 2 | N/A [a] | **217±13** | 204±32 [c] | **450±17** | **313±17** |
| | 5 | N/A [a] | 114±92 [a] | 76±104 [a] | **246±16** | - |
| | 10 | - | N/A [a] | N/A [a] | 127±109 [a] | N/A [a] |
| 150 | 1 | N/A [a] | **368±14** | **341±25** | - | N/A [b] |
| | 2 | N/A [a] | **243±10** | **254±16** | **473±14** | **355±5** |
| | 5 | N/A [a] | 131±108 [a] | N/A [a] | 264±15 [c] | 213±8 [c] |
| | 10 | - | N/A [a] | N/A [a] | 131±122 [a] | N/A [a] |
| 180 | 1 | 337±50 [c] | - | - | - | - |
| 200 | 1 | N/A [a] | **418±13** | **404±21** | - | N/A [b] |
| | 2 | N/A [a] | **273±10** | 293±19 [c] | N/A [b] | **412±8** |
| | 5 | N/A [a] | 134±61 [c] | N/A [a] | **296±21** | **246±13** |
| | 10 | N/A [a] | N/A [a] | N/A [a] | 202±107 [a] | N/A [a] |
| 250 | 1 | 676±133 [b] | **421±17** | **494±26** | - | - |
| | 2 | 303±65 [c] | **292±13** | 324±41 [c] | N/A [b] | **440±18** |
| | 5 | 293±59 [a] | 157±66 [c] | 169±125 [a] | 381±59 [c] | **264±15** |
| | 10 | 159±79 [a] | N/A [a] | N/A [a] | 319±37 [c] | N/A [a] |
| 300 | 1 | N/A [b] | **590±10** | - | - | - |

| | | | | | | |
|---|---|---|---|---|---|---|
| | 2 | - | **364±15** | - | - | N/A [b] |
| | 5 | 604±127 [c] | 214±89 [c] | - | N/A [b] | **337±12** |
| | 10 | 337±51 [c] | N/A [a] | - | 367±38 [c] | 244±36 [c] |
| 350 | 1 | - | **652±21** | N/A [b] | - | - |
| | 2 | - | **396±11** | 492±56 [b] | - | N/A [b] |
| | 5 | - | 271±18 [c] | 294±113 [a] | N/A [b] | 358±10 [c] |
| | 10 | 569±85 [c] | 181±29 [c] | N/A [a] | 368±26 [c] | 243±53 [c] |
| 400 | 1 | - | **622±16** | N/A [b] | - | - |
| | 2 | - | **412±18** | N/A [b] | - | - |
| | 5 | - | 274±9 [c] | 357±62 [c] | N/A [b] | 440±11 [c] |
| | 10 | - | 168±33 [c] | 230±155 [a] | 444±15 [b] | 294±20 [c] |
| 450 | 1 | - | **762±16** | - | - | - |
| | 2 | - | **480±21** | N/A [b] | - | - |
| | 5 | - | 258±19 [c] | 364±42 [c] | - | **466±16** |
| | 10 | - | 205±53 [c] | 280±113 [a] | 456±17 [b] | 301±27 [c] |
| 500 | 1 | - | - | **-** | - | - |
| | 2 | - | 563±17 [c] | N/A [b] | - | - |
| | 5 | - | 308±19 [c] | 405±32 [c] | - | N/A [b] |
| | 10 | - | **243±21** | 260±102 [a] | 458±27 [c] | 302±36 [c] |
| 550 | 5 | - | - | 422±87 [b] | - | - |
| | 10 | - | - | 261±101 [a] | 461±14 [c] | 380±21 [c] |
| 600 | 5 | - | - | 447±53 [c] | - | - |
| | 10 | - | - | 254±79 [a] | N/A [b] | 393±25 [c] |
| 650 | 5 | - | - | 481±58 [b] | - | - |
| | 10 | - | - | 340±63 [c] | N/A [b] | 446±70 [b] |
| 700 | 5 | - | - | N/A [b] | - | - |
| | 10 | - | - | 288±58 [c] | N/A [b] | N/A [b] |

N/A - not applicable, single strand could not be measured; a - no material was extruded or broken strand was printed (pressure not sufficient or speed too high); b - strands merged (pressure too high), c - strands not smooth, with defected edges; " –" printing not attempted. Data obtained for smooth (no failure in the edges) and continuous, not interrupted strands are marked in bold.

### 5) Printing in 2D of PEG-Dop/$Al^{3+}$ (pH 8.4) and PEG-Dop/$Fe^{3+}$ (20 kDa) inks

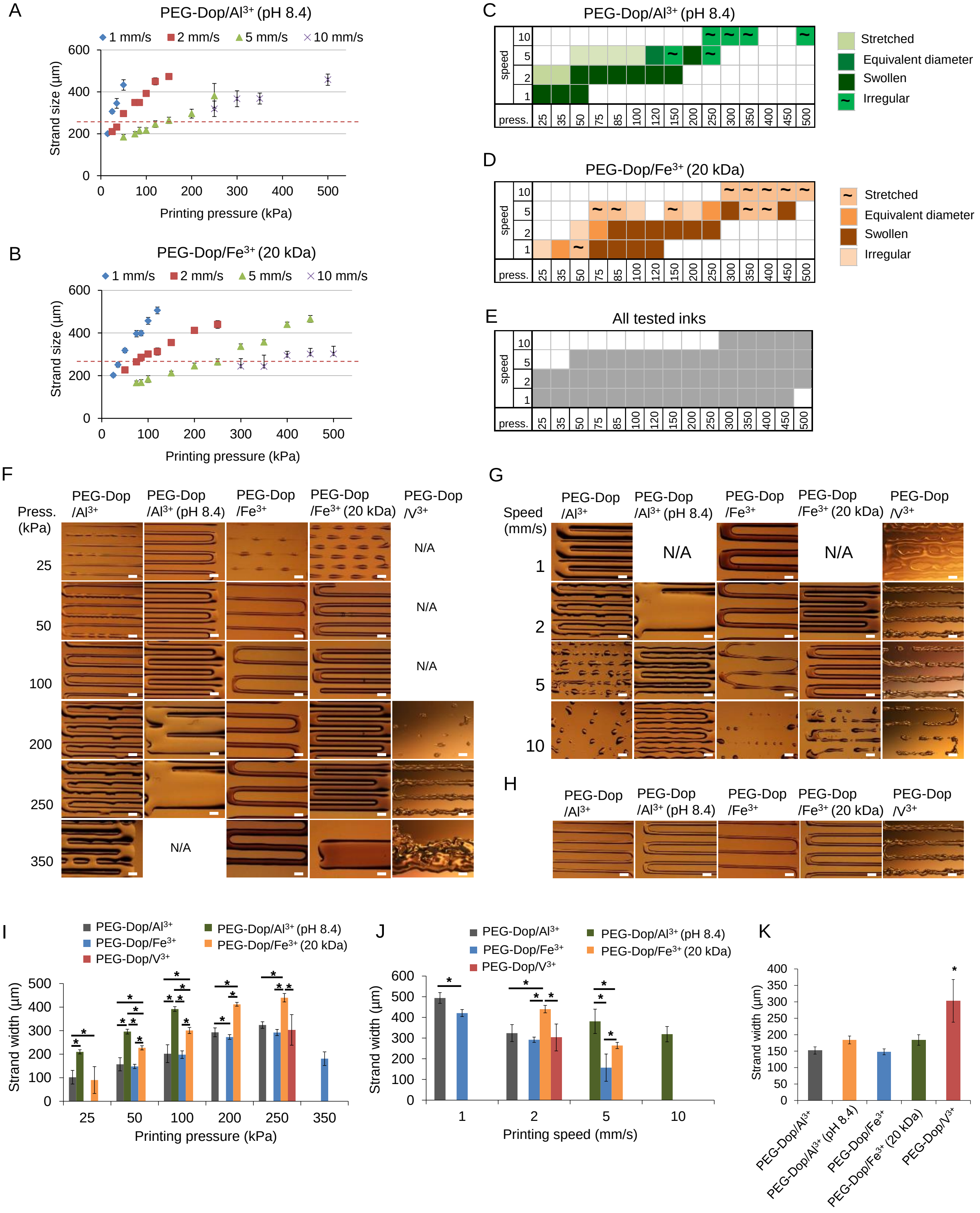

**Figure.S6.** Printability of the tested inks. Strands diameter depending on head moving speed (mm/s) and printing pressure (kPa) for: PEG-Dop/$Al^{3+}$ (pH 8.4) (**A**) and PEG-Dop/$Fe^{3+}$ (20 kDa) (**B**). Red lines serve as a guide for the eye showing the strand width value equal to the printing nozzle diameter (260 µm). Printability range for PEG-Dop/$Al^{3+}$ (pH 8.4) ink (**C**), PEG-Dop/$Fe^{3+}$ (20 kDa) ink (**D**) and all 5 inks tested in the study (**E**). Influence of pressure on printing of different inks at the head moving speed of 2 mm/s (**F**). Influence of head moving speed on printing of different inks at pressure of 250 kPa (**G**). Printed lines with the highest accuracy, defined as the thinnest printed strands with possibly smooth edges (**H**). Quantification of strand diameters corresponding to C, D, E, (**I**, **J**, **K**, respectively). Scale bars: 500 µm.

6) Printing in 3D

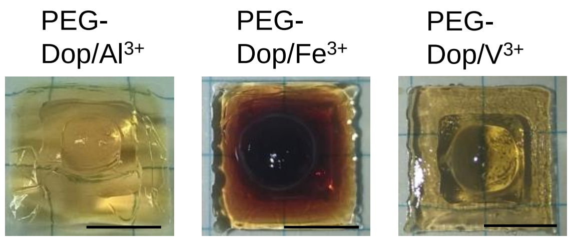


**Figure S7.** Printing of bulk 3D scaffolds with PEG-Dop/$V^{3+}$, PEG-Dop/$Fe^{3+}$ and PEG-Dop/$Al^{3+}$ inks. Pyramid-like printed constructs were composed of 3 levels, 4-5 layers each, with dimensions: 1$^{st}$ level 9.1 mm x 9.1 mm, 2$^{nd}$ level 6 mm x 6 mm and 3$^{rd}$ level 3 mm x 3 mm. Parallel strands forming each layer were printed in contact to form bulk structure (strand distance: 0,6 cm – 1cm). Each level of construct was crosslinked with 2 ml of crosslinking solution for 30 seconds, before printing of the following level. Printing was performed at 1 mm/s speed, and 120 kPa pressure for PEG-Dop/$Al^{3+}$ and PEG-Dop/$Fe^{3+}$, 200 kPa pressure for PEG-Dop/$V^{3+}$. The z-offset for printing of consecutive levels was chosen while printing based on the estimated height of the previously printed and crosslinked level. Scale bars: 500 µm.

7) Cell viability studies for PEG-Dop/$Al^{3+}$ (pH 8.4) and PEG-Dop/$Fe^{3+}$ (20 kDa)

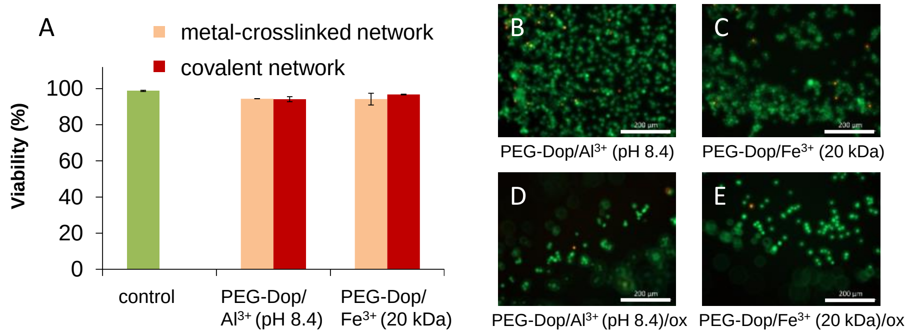


**Figure S8.** Fibroblasts viability after 1h culture in direct contact with PEG-Dop based networks. **A.** Cell viability in contact with metal-crosslinked network (orange bars) and covalently crosslinked network (red bars), in comparison to cells cultured on plastic (green bar). No significant differences were found. Life/dead staining exemplary images of fibroblasts cultured with metal-crosslinked networks (**B**: PEG-Dop/$Al^{3+}$ (pH 8.4), **C**: PEG-Dop/$Fe^{3+}$ (20 kDa)) and covalent networks crosslinked with oxidant (ox) (**D**: PEG-Dop/$Al^{3+}$ (pH 8.4), **E**: PEG-Dop/$Fe^{3+}$ (20 kDa)). Scale bars: 200 µm.

**References:**

[1] H. Li, S. Liu, L. Lin, Rheological study on 3D printability of alginate hydrogel and effect of graphene oxide, International Journal of Bioprinting 2(2) (2016) 54-66.
[2] R. Suntornnond, E.Y.S. Tan, J. An, C.K. Chua, A Mathematical Model on the Resolution of Extrusion Bioprinting for the Development of New Bioinks, Materials (Basel), 2016.
